# CagY sequence and structural motifs are associated with ancestry and disease in world-wide *Helicobacter pylori* strains

**DOI:** 10.1101/2025.06.20.660796

**Authors:** Mario Angel López-Luis, Roberto C Torres, Jay V. Solnick, M. Constanza Camargo, Alfonso Méndez-Tenorio, Javier Torres

**Affiliations:** Departamento de Bioquímica, Laboratorio de Biotecnología y Bioinformática Genómica, Escuela Nacional de Ciencias Biológicas, Instituto Politécnico Nacional, Campus Lázaro Cárdenas; Unidad de Investigación en Enfermedades Infecciosas, UMAE Pediatría, Instituto Mexicano del Seguro Social, Ciudad de México, México; Department of Medicine and Department of Microbiology and Immunology, University of California, Davis, CA, USA; Division of Cancer Epidemiology and Genetics, National Cancer Institute, Rockville, MD, USA

## Abstract

*Helicobacter pylori* (*Hp*) interacts with gastric epithelial cells using a multi-protein type 4 secretion system (T4SS) whose function is regulated by CagY. CagY is the T4SS backbone, connecting the bacterial cytoplasm with the epithelial cytoplasm and interacting with most other T4SS proteins. However, its mechanisms of action are unknown. We aimed to comprehensively analyse *cagY* in a worldwide collection of strains to unmask any correlation between ancestry and/or disease and sequence diversity, including an analysis of critical domains and associated structural changes. We used 674 *cagY* sequences derived from1,012 *Hp* genomes, sequenced with Single Molecule Real-Time from the *Hp* Genome Project. Phylogenetic and principal component analyses (PCA) revealed a strong population structure among European, Asian, African and American clusters, which interfered with other analyses. To address this heterogeneity, dimensional reduction analysis with PCA and Linear Discriminant Analyses were used to minimize interference before studying possible association of *cagY* diversity with disease. For this analysis, *cagY* genes were fragmented using unitig-caller with a designed analytical pipeline that clearly separated the non-atrophic gastritis (NAG), advanced intestinal metaplasia and gastric cancer (GC) groups. Sixty-four unitigs significantly separated GC from NAG based on *cagY* sequences, and the most GC-associated unitigs were localized in the middle repeat region of the protein (MRR), particularly in the A module, including the cysteine-containing motif YLDCVSQ. Proline was enriched in the MRR and VirB10 regions. The consequences of the diversity on the protein structure were analysed using ChimeraX. The validated and stabilized 14-multimer model showed major structural changes along the multimer-structure, particularly in the cysteine-rich MRR region, that might be one mechanism by which CagY regulates T4SS function.

## Introduction

*Helicobacter pylori* (*Hp*) colonizes the gastric epithelium of about 43% of the human world population, and it is usually acquired in early childhood due to intimate contact with mother and other family members infected with the bacteria (Li et al., 2023). The infection is asymptomatic in most individuals, although about 2% of those infected will develop gastric cancer (GC) after decades of gastric colonization (Herrera & Parsonnet, 2009) . *Hp* has coevolved with humans most probably since the origin of our species (Falush et al., 2003), resulting in an exquisite adaption to live in the hostile environment of the stomach. Accordingly, it is not a surprise to learn that the population structure of *Hp* mirrors that of humans and can be used as an excellent marker of ancient and recent human migrations (Falush et al., 2003). In addition, the ancestry of *Hp* strains may be studied with fineSTRUCTURE, a program developed to study human populations (Moodley et al., 2009).

The best documented *Hp* virulence factor is the pathogenicity island, cagPAI, consisting of operons with about 17 genes that codes for a T4SS, a complex structure that spans from the *Hp* cytoplasm through the bacterial membrane and facing out to interact with the membrane of the gastric epithelial cell. This T4SS allows the translocation of DNA, bacterial cell envelope precursor metabolites and the oncogenic protein CagA to the gastric epithelial cells. Once in the cytoplasm, the EPIYA motifs of CagA become phosphorylated in order to interact with kinases and other cellular molecules, leading to activation of inflammatory mediators and cytoskeletal modifications (Alipour, 2029; Sharafutdinov et al., 2021), increasing the risk for the development of gastric diseases.

CagY is a key protein for the function of the T4SS, regulating its pro-inflammatory activity, modulating the intensity of gastric inflammation and hence the tissue damage (Delahay et al., 2008). CagY is a 2,000 aa protein with an extremely unusual high number of cysteines (around 58 residues) and it is one of the largest and more complex bacterial proteins. Its sequence contains different domains that function as the tunnel of the T4SS “syringe”, connecting the cytoplasm of Hp with the cytoplasm of the epithelial cell, spanning the bacterial internal membrane (IM), periplasm and the outer membrane (OM), to fa+-ce outside (Hu et al., 2019). Furthermore, CagY includes two polymorphic repeated regions, the Five Repeat Region (FRR) an intrinsically disorder region (IDR) with unknown function and the Middle Repeat Region (MRR) an 800 amino acids domain with two repetitive modules, A and B. Each module includes the motifs: delta (δ), mu (μ) and alpha (α) for module A and epsilon ((ε), lambda (λ) and beta (β) for module B (Delahay et al., 2008; Liu et al., 1999). In addition, it has the antenna projection (AP), a region located at the tip of the T4SS that interacts with the gastric epithelial cells (Tran et al., 2023). It is important to note that CagY does not share homology with any known protein, except for a short region in the carboxyl terminus, homologous to VirB10 (Barrozo et al., 2016; Delahay et al., 2008; Tran et al., 2023). Furthermore, several of the other Cag proteins bind along the different domains of CagY to form the complex T4SS (Backert & Tegtmeyer, 2017; Delahay et al., 2008; Fischer et al., 2020). It is not a surprise then that CagY seems to have the key function of modulating the activity of the T4SS working as a rheostat to turn on and off the adaptative immune response (Barrozo et al., 2016; Sierra et al., 2019; Tegtmeyer et al., 2020). The mechanism to regulate this process is uncertain, although there is evidence suggesting that recombination between or mutations within the repeat modules in the MRR region are associated with the rheostat function (Barrozo et al., 2013, 2016; Delahay et al., 2008; Sierra et al., 2019). In addition, it has been suggested that CagY may also induce inflammation by interacting with TLR5, to produce Interleukin 8 (IL8) (Tegtmeyer et al., 2020).The VirB10 homologous region is also reported to bind to integrins in the outer membrane of gastric epithelium cells, an activity that may depend on the composition of the MRR region (Koelblen et al., 2017).

Despite its importance for the function and virulence of the T4SS, very little is known about the mechanisms of action and about structure-function of CagY. The study of this protein is very challenging because of its high sequence diversity, multiple domains, large size including the 800 amino acids repeats region and its multimer structure (Delahay et al., 2008). Furthermore, because of its highly diverse gene sequence and unusually long repeats region, the sequences deposited in databases are often not reliable because they are usually truncated or incorrectly annotated (Liu et al., 1999). Only a fraction of the protein has been crystalized and its structure as a monomer or multimer is yet unknown (Sheedlo et al., 2020). We were recently able to model the structure of the CagY multimer using several different approaches and considering the reported crystalized regions of the protein (López-Luis et al., 2023). In this work we aimed to comprehensively analyse *cagY* and its encoded protein to learn about the correlation of sequence-structure-function-disease. For this analysis, we used the genomes from the *H. pylori* Genome Project (*Hp*GP) (Thorell et al., 2023), the largest worldwide collection of strains sequenced using long-read sequencing. The *cagY* sequences were analysed with an array of machine learning methods, because of their ability to discern complex patterns or to make high confidence modelling simulations (Jumper et al., 2021) plus other recently developed bioinformatic tools, including by ours (López-Luis et al., 2023).

## Methods and material

### *Helicobacter pylori* dataset

The *Hp*GP was launched by the US National Cancer institute (*Hp*GP, NCI, USA) and the *H. pylori* strains were sequenced with Single Molecule Real-Time (SMRT)/Pacific Biosciences (PacBio) (Eid et al., 2009). A total of 1,012 *H. pylori* genomes were provided by the NCI with metadata including ancestry of the strains and clinical condition of the individuals (Thorell et al., 2023). The genomes were filtered for the presence of cagPAI and then the *cagY* gene was extracted from Prokka v1.14.6 (Seemann, 2014) and HMMer v3.2.1 (Wheeler & Eddy, 2013) outputs, extracting 1000 nucleotides upstream and downstream of *cagY*, to avoid truncated sequences. For validation of results nine previously reported *cag*Y sequences were included in the analyses (Barrozo et al., 2013; Jackson et al., 2020). We finally studied 682 sequences of *cagY* from different countries (Suppl Table 1).

### Phylogenetic analysis of *cagY* gene

The *cagY* gene was aligned using a Hidden Markov Model (HMM) with the 100 longest *cagY* genes, constructed with HMMer v3.2.1 software (Wheeler & Eddy, 2013) followed by a progressive alignment of all *cagY* genes with ClustalW v2 (Larkin et al., 2007). Then, a phylogenetic tree was constructed with Maximum Likelihood using GTR model followed by independent rate of inversions and transversions with MEGA v11 Software (Tamura et al., 2021) and visualized with itol v7.2 (Letunic & Bork, 2024). The Single Nucleotide Polymorphisms (SNPs) were extracted from the *cagY* alignment with SNP-sites v2.4.1 (Page et al., 2016) and a Principal Component Analysis (PCA) of the presence/absence of SNPs constructed with a script in python using the scikit-learn library (Pedregosa et al., 2011) and RStudio and finally plotted with ggplot2 library (Wickham, 2016).

### Estimation of Virtual Genomic fingerprints with VAMPhyRE

Virtual Genomic Fingerprints (VGFs) were built with Virtual Analysis Method for Phylogenomic fingeRprint Estimation (VAMPhyRE) using a library of 65,536 probes of 8-mers, as previously described (Muñoz-Ramírez et al., 2017). The obtained table of phylogenetic distances was analyzed with MEGA v11 (Tamura et al., 2021) using Unweighted Pair Group Method with Arithmetic Mean (UPGMA) and itol v7.2 (Letunic & Bork, 2024). In addition, a dimensional reduction analysis was done with PCA and informative features with an importance value (Mean Decrease Gini, MDG) >zero selected using RandomForest (RF) and displayed by ggplot2 library (Wickham, 2016) in R.

### Phylogenetic network of *cagY* gene

Genetic distances between each pair of strains were obtained with Nexus and formatted as a triangular distance matrix, as previously described (Muñoz-Ramirez et al., 2021). Each distance was normalized between 0 and 1 according to:

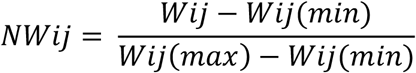

where NWij is the normalized weights between strains i and j, Wij is the genetic distance between strains i and j, and Wij(max), Wij(min) are the maximum and minimum value for the genetic distance in the whole network respectively. With this normalization a value of 0 means the two strains have the highest genetic similarity, and a value of 1 means the two strains are the most dissimilar.

### Analyses of *cagY* variants and their probable association with disease

To evaluate the association between *cagY* diversity and gastric diseases, we implemented the fragmentation of *cagY* gene into unitigs using unitig-caller v1.3.1 (Holley & Melsted, 2020). The program constructs a database of unique unitigs with length of at least 31 pb in each *cagY* sequence, and a presence/absence matrix was built for each of the 674 *cagY* sequences with a python script. Next, a dimensional reduction analysis with PCA and LDA was run. For the LDA analyses we selected the most informative unitigs with MDG values >zero using Random Forest (RF). For the PCA we used the first 20 principal components (PC) with all unitigs and select the more informative for each PC components. For comparation of the presence/absence of unitigs in GC vs NAG and IM vs NAG a Fisher Exact Test was used, and the p-value corrected with Bonferroni. Finally, we aligned the unitigs to *cagY* of the reference strain 26695-ATCC of *Hp* to localize the significant unitigs (p-value <0.01) using a Python script. The results were represented in a Manhattan plot, illustrating the unitig position in *cagY* and its p-value in each disease group.

### Classification and construction of Machine Learning model

To train and evaluate the prediction capability of Machine Learning (ML) models for disease, a frequency matrix with all unitigs was constructed (including those that significantly distinguished the groups) with a pipeline developed in Python v3.9 and scikit-learn library (Pedregosa et al., 2011). The following models were tested, Multinomial Naive Bayes (MNB), Support Vectorial Machine (SVM), Logistic Regression (LR), RandomForest (RF) and Multi-Layer Perceptron (MLP) as simple Neural Network (NN). For RF method we determined the importance of each unitig with the MDG value. All classification algorithms were used with default values. The performance of informative unitigs was estimated using ROC-AUC and F1 Score used for training and 20% to test data, samples we paired using the Random over sampler of scikit learn, and a 10-training with different seed was done (Asnicar et al., 2024; Mahesh, 2020; Raimondi et al., 2019). All methods were compared with Wilcoxon Rank Sums Test and a p-value < 0.05 was considered as significant.

### Localization and co-presence of informative unitigs along CagY

To investigate whether informative unitigs exhibit disease-specific co-occurrence patterns suggestive of distinct secondary-structural arrangements, we used the *cagY* sequence of reference strain 26695 as a scaffold for unitig mapping. All informative unitigs were aligned to the 26695 loci via a custom Python pipeline. We then assessed the pairwise presence/absence of variants across the gene using Fisher’s exact test, and plotted significative co-occurrence events, together with their genomic coordinates and association p-values, in a Manhattan plot generated with ggplot2 (Wickham, 2016) in RStudio.

### Homology modelling and Molecular dynamics of CagY

The effect of the sequence variation on the structure of *cagY* was estimated doing a homology modeling of the CagY protein with MODELLER (v10.4) (Eswar et al., 2006), with python 3.9, using CagY of *Hp* 26695 as a reference, as previously described (López-Luis et al., 2023). 1000 homology models were constructed for each CagY sequence to select the best model based on the DOPE score. The tridimensional model was aligned with ChimeraX software (Pettersen et al., 2021) based on CagY 26695 topology. To stabilize and refine the model, we constructed a trimmer of each CagY protein, and the system of explicit solvent was constructed with VMD software (Humphrey et al., 1996), using the recommended script with NAMD3 software (Phillips et al., 2020) for 10 to 50 ns. The Root Mean Square Deviation (RMSD) and Root Mean Square Fluctuation (RMSF) along Equilibrium Molecular Dynamics (EMD) were calculated with Python MDAnalysis (Cohen et al., 2023). Finally, the comparation of position of RMSD between CagY structures was made with ChimeraX. The plots were constructed and displayed with ggplot2 in Rstudio.

## Results

### Phylogenetic analyses reveal a strong population structure for *cagY* gene and for the MRR region of the gene

Phylogenetic reconstruction of *cagY* using a kmer based reconstruction (Figure 1a). showed a strong population structure with clusters for European, Asian, African and Latin American populations, similar to what we and others have previously observed with the analyses of the core genome (Muñoz-Ramírez et al., 2017; Yahara et al., 2013) suggesting a strong coevolution of *cagY* enough to mirror the ancestry of its human host. Another approach, a phylogenetic network analyses also confirmed the strong population structure of *cagY* (Figure 1b), depicting the grouping of the gene sequences following their ancestry. Even the analyses of only the MRR region showed a population structure like the whole *cagY* gene (Figure 1c). Furthermore, an LDA supervised reduction dimension of *cagY* and MRR sequences also showed a clear separation following ancestry (Suppl Figure 1a and 1b). Thus, phylogenetic and reduction dimension approaches confirmed the strong population structure of *cagY* and its MRR fragment.

**Figure 1.**
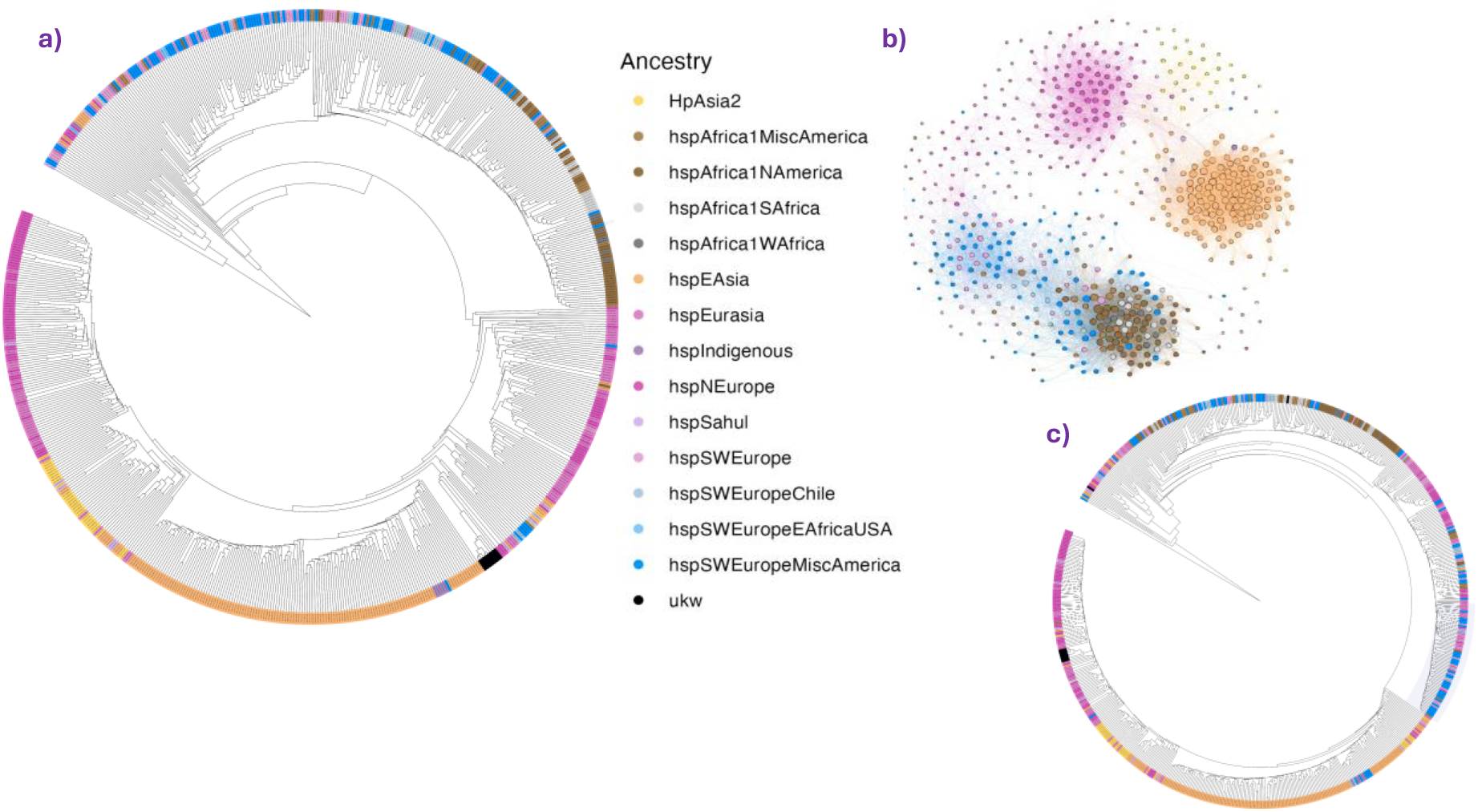
Phylogenetic analysis of the c*agY* gene using the VAMPhyRE strategy. Colours represent the ancestry of *Helicobacter pylori*. **a)** Phylogenetic analysis of the complete *cagY* gene, showing the clustering of *H. pylori* ancestries, which mirrors the clustering of the core genome. **b)** The tree cluster related *Hp* populations, distinguishing the ancestries of Asia-Europe and the America-Africa regions. **c)** Phylogenetic analysis of the MRR region of the *cagY* gene, showing the ancestry clustering of the MRR with lower resolution compared to the complete *cagY* gene.

### Bioinformatic tools using unitigs reveal association of *cagY* diversity with disease

Once the strong population structure of *cagY* was revealed, the challenge was to avoid its effect that confounds other analyses and search for any correlation of *cagY* variants with disease. After trying several tests, we found an approach that could distinguish the clinical groups, the fragmentation of each of the *cagY* genes into unitigs to construct a library and scan back all genes, a strategy previously used to more efficiently compare genomes in Genome Wide Association Studies (GWAS). After scanning each gene with the unitigs library, a presence/absence matrix was built to run a dimensional reduction analysis with PCA (Suppl Figure 2) and as expected, components C1 and C2 clearly clustered following the *Hp* ancestries (Suppl Figure 2a), but not the clinical source of the *cagY* sequence (Suppl Figure 2b). A variance analysis showed that the first 20 components explained almost 60% of the variance. According to these results another dimensional reduction analysis using a supervised LDA was implemented, to detect “hidden” patterns associated with disease. The resulted LDA plot showed a clear discrimination of the clinical groups, NAG, IM and GC (Figure 2a), LD1 discriminated GC from IM, whereas LD2 discriminate NAG from GC and IM. We then asked to which components of the PCA analysis (Suppl Table 2) were associated the unitigs most informative to discriminate the disease group and for that we determined the mean decrease gini value (MDG) of each informative unitig (Figure 2b). Most of the unitigs with the highest MDG value were associated with components 10 or higher, and although some were also informative for ancestry (PC1-associated) it was possible to identify unitigs informative for disease but not for population structure. These results indicate that our pipeline analyses made a correct adjustment for population structure, resulting in a reliable approach to predict the clinical origin based on *cagY* diversity.

**Figure 2.**
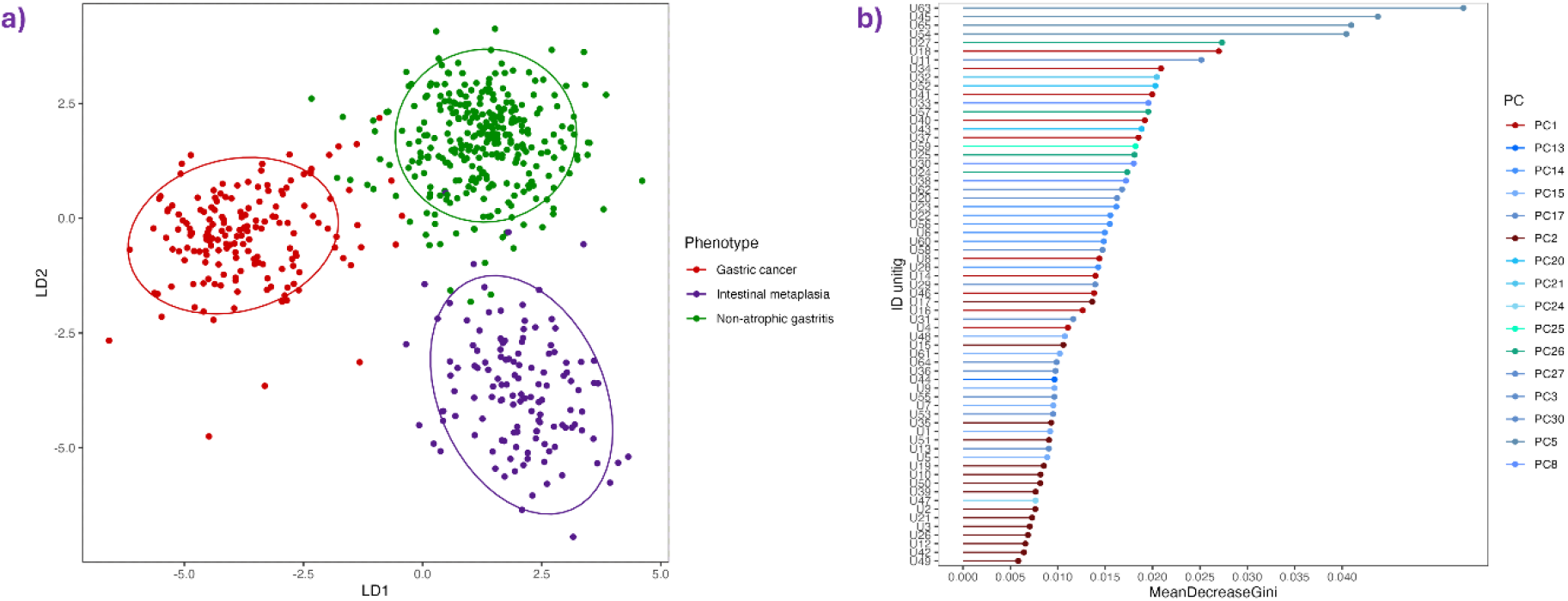
Machine Learning analysis of *cagY* unitigs. **a)** Linear Discriminant Analysis (LDA) of the Presence/Absence matrix of unitigs. The analysis shows the phenotype-based clustering. **b)** Mean Decrease Gini (MDG) values of significant unitigs, where the colour represents the Principal Component (PC) associated with each unitig. The highest MDG values are related to higher PCs, while the lowest values are associated with PC 1 and PC 2.

### Short fragments of *cagY* (unitigs) may distinguish variants of *cagY* associated with GC

The unitigs significantly associated with each clinical group were identified and aligned to CagY from *Hp* 26695 reference strain (Suppl. Table 2). Sixty-five unitigs were significantly associated with GC, whereas no unitig was found significantly associated with IM. Accordingly, the group of IM was not included in the following analyses, which was performed using NAG as controls and GC as cases. The ability of the 65 GC-associated unitigs to predict disease group was validated with supervised ML methods. Five supervised ML models were trained using three set of unitigs, all 22,721 unitigs from the generated library, the 65 GC-associated unitigs and 65 randomly selected unitigs (Suppl Table 3). The RF model was the best prediction model with a ROC-AUC value over 83% and a F1 Score of 80% with the 65 GC-associated unitigs. In contrast, the analyzes with all-library unitigs, resulted in significantly lower ROC-AUC values and F1 Scores (Suppl Figure 3).

### Sequence of the unitigs informative for GC locate mostly in the MRR of CagY

Once the informative value of the GC-associated unitigs was validated, their position was located by aligning the sequences in the CagY protein of Hp-26695 reference strain (Figure 3a) and the functional domain identified. The 65 unitigs were identified in 79 different positions, 65 were informative for GC and the other 14 for NAG (Suppl Table 4). Most of the GC-associated unitigs located at the MRR region and 13 mapped repeatedly in 37 different positions, starting at the 8^th^ repetitive module corresponding to an A module. Among the 37 unitigs, 32 mapped in the A module mostly in the delta-mu motifs, and only 5 located in the B module. The predominant location of GC-associated unitigs at the MRR is in accordance with previous reports that have associated this region with modulation of the function of the T4SS (Barrozo et al., 2013, 2016; Bella et al., 2021; Delahay et al., 2008; Liu et al., 1999; Tegtmeyer et al., 2020). Additionally, we found the presence of cysteine codifying codons in 25 of the 37 unitigs mapping within MRR (Figure 3a), an important observation because of the role of these many cysteine residues in the structure of the protein and its final multimer, as described below. Other informative unitigs located outside the MRR, mainly in the VirB10 region and between MRR and VirB10 (Figure 3a). All the NAG-associated unitigs mapped outside MRR, between MRR and VirB10 and in the FRR domain. In addition, we performed an analysis of copresence of pairs of unitigs, to identify those that were significantly co-present in the GC group, as illustrated in the Manhattan plot of Figure 3a. The analyses showed that some unitigs in the MRR region were significantly co-present with unitigs outside MRR, particularly in the VirB10 domain.

**Figure 3.**
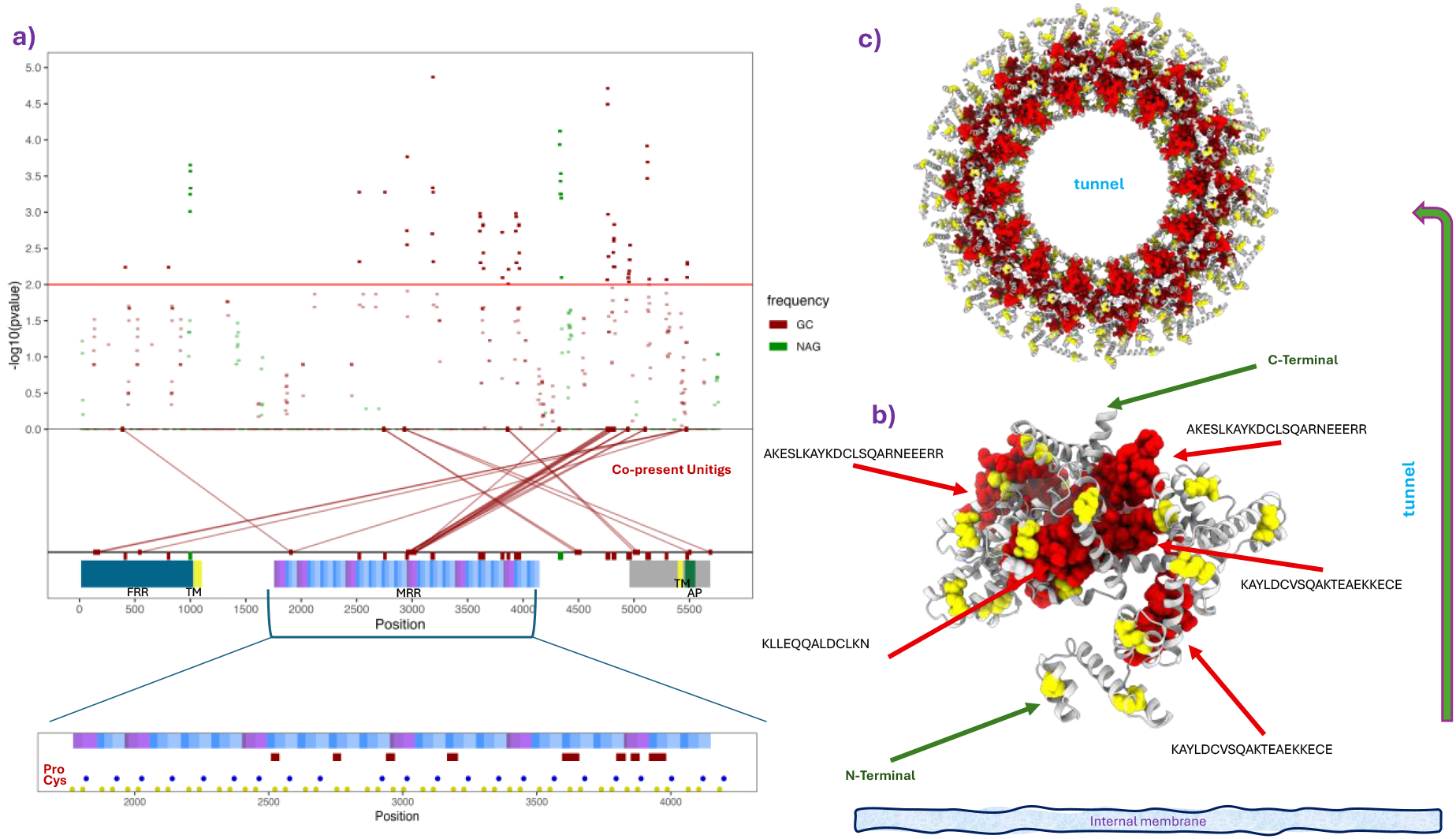
Unitig association analysis of the *cagY* gene. **a)** Manhattan plot of unitig-phenotype associations, where colours represent the frequency in each phenotype, green to NAG and Red to GC. Below the plot, the copresence analysis, functional domains, and cysteine and proline codons positions are displayed. The analysis highlights significant associated regions in *cagY* represented as unitigs. The copresence analysis illustrates intragenic GC interactions. **b)** Three-dimensional structure of the MRR region, showing the representation of unitigs and cysteine positions. **c)** Three-dimensional structure of the 14 subunits of the MRR region, illustrating the presence of unitigs and cysteine positions.

### Tridimensional structure of CagY-MRR highlights the importance of associated unitigs

Once the informative unitigs were mapped in the amino acid sequence of CagY we moved to elucidate the effects these variants may have on the tertiary structure of the protein using the analytical pipeline we recently developed to model CagY (López-Luis et al., 2023). We first mapped the GC-informative unitigs in the structure of MRR, where many included a Cys residue as described before (Figure 3a). In the monomer the unitigs concentrated in a highly folded Cys-rich region sitting in the periplasm of the membrane (Figure 3b). The multimer (14-mer) of the protein forms a donut at the base of the T4SS with the MRR region, which lumen is supposed to be the channel through which CagA and other molecules are translocated (Figure 3c). The figure shows that unitigs including cysteine form concentric rings shaping the donut where the AYLDCVSQAK uniting (U27) located in the inner ring, facing to the lumen of the tunnel. The composition of this structure suggests that variation in the aa sequences around the Cys bridges could result in modification of the diameter of the tunnel. To further study this possibility, we next analyzed the structure of the 14-multimer and began with the reference HP26695 CagY to spot the known domains and the location of the informative unitigs (Figure 4a). Whereas most of the GC-associated unitigs located at the base on the amino side of the protein, some also localized up in the neck and in the antenna region (AP) within the VirB10 homologue motif. Figure 4b shows the localization of the informative unitigs in the 14-multimer, where it is clear how the unitigs interact to form rings along the complex CagY-multimer that constitute the skeleton of the T4SS.

**Figure 4.**
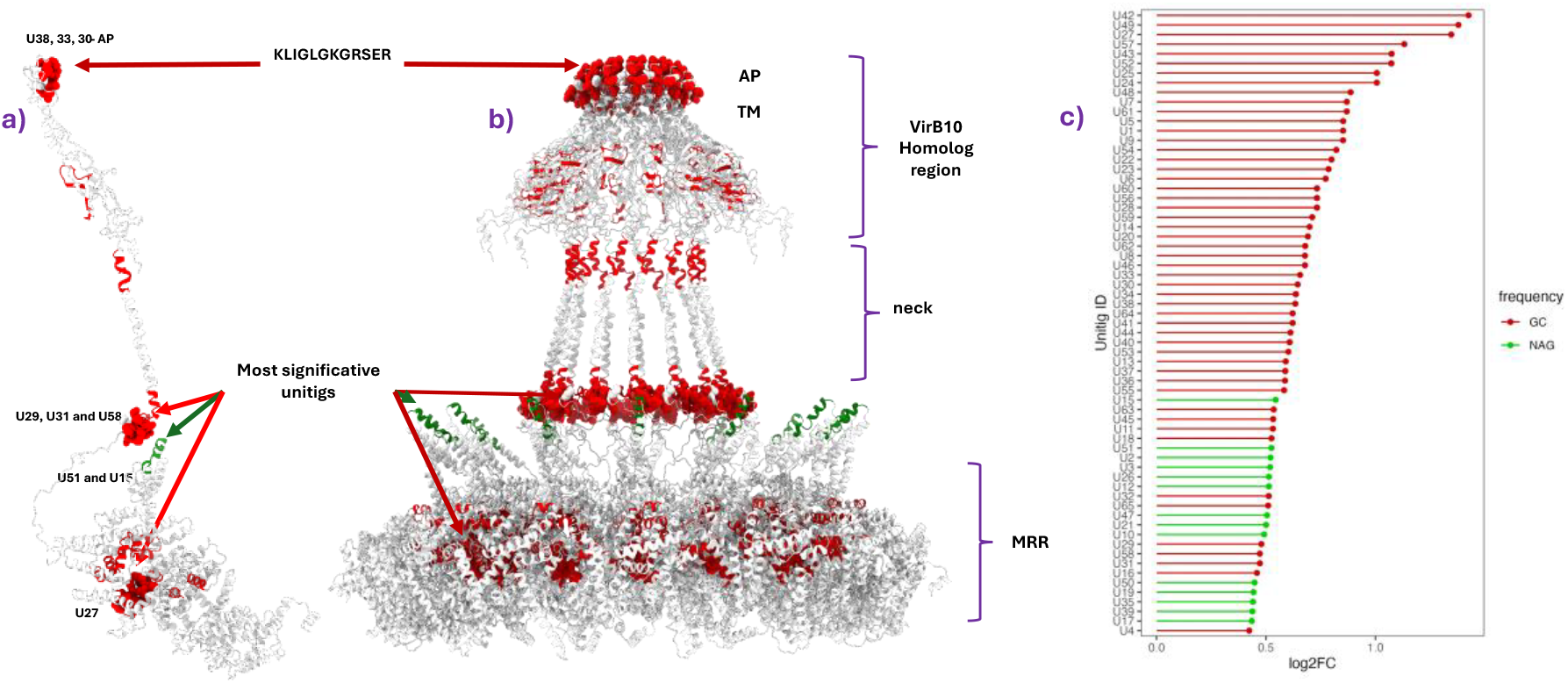
Three-dimensional representation of unitigs in CagY 26695. Purple represents the MRR region, dark gray represents the VirB10 homologous region, yellow represents the TM region, red and green represent the association of unitigs, and orange represents cysteine positions. **a)** Three-dimensional structure of the CagY multimer, showing the representation of unitigs within the CagY multimer. **b)** Monomer structure of CagY, in this structure we see the unitigs amog CagY sequence. **c)** log2 of Fold Change of unitig presence whitin CagY phenotypes.

Once the model with the reference sequence was ready, the analyses was done with sequence variants associated to different diseases but with similar ancestry (hspSWEuropeMiscAmerica) two NAG, two IM and two GC. The stabilized models of the six CagY sequences (Suppl. Figure 4a) were different from each other, as evidenced by the RMSD values of comparing structures from the same disease (Table 1), even when we compare the structures of same phenotype (Suppl. Figure 5). Validation analyses for the six models are shown in suppl Figure 5a. The stabilized and validated 14-mer model for the six proteins is presented in Figure 5, where marked differences between proteins can be observed all along the multimer, from the antenna and down to the MRR region. In the above row of figure 5, CagY from the NAG strain HpGP-COL-120 shows the lumen of the tunnel closed, whereas in IM (strain HpGP-MEX-001) and GC (strain HpGP-PER-114) the tunnel is open. However, in the row below, all three multimers from NAG, IM and GC, respectively, present the tunnel open. Among the many changes, we noticed that the structure of the two GC sequences had a narrower diameter in the neck region (orange arrow in Figure 5). It seems that the structure of the CagY multimer is as diverse as the sequences, and it is hard to identify any changes associated with the disease of origin.

**Table 1.** Determination of the Root Mean Square Deviation value comparating between each of the eight CagY structures analyzed.

| Strain | HpGP-PER-136 | HpGP-PER-114 | HpGP-MEX-006 | HpGP-MEX-001 | HpGP-COL-119 | HpGP-COL-120 | rOut1-<br>IL8- | rOut3_IL<br>8+ | J166_IL8<br>+ |
| --- | --- | --- | --- | --- | --- | --- | --- | --- | --- |
| HpGP-PER-136-GC | — | 35.924 | 15.797 | 38.51 | 19.438 | 19.69 | 25.96 | 22.96 | 29.05 |
| HpGP-PER-114-GC | 35.924 | — | 48.472 | 22.414 | 32.646 | 31.724 | 44.45 | 30.92 | 39.2 |
| HpGP-MEX-006-NAG | 15.797 | 48.472 | — | 32.82 | 19.629 | 16.514 | 20.42 | 18.86 | 23.5 |
| HpGP-MEX-001_IM | 38.51 | 22.414 | 32.82 | — | 35.637 | 31.438 | 42.2 | 27.86 | 46 |
| HpGP-COL-119_IM | 19.438 | 32.646 | 19.629 | 35.637 | — | 14.296 | 25.95 | 14.73 | 22.22 |
| HpGP-COL-120-NAG | 19.69 | 31.724 | 16.514 | 31.438 | 14.296 | — | 19.03 | 13.66 | 23.7 |
| rOut1-IL8- | 25.96 | 44.45 | 20.42 | 42.2 | 25.95 | 19.03 | — | 25.66 | 18.51 |
| rOut3-IL8+ | 22.96 | 30.92 | 18.86 | 27.86 | 14.73 | 13.66 | 25.66 | — | 20.19 |
| J166_IL8+ | 29.05 | 39.2 | 23.5 | 46 | 22.22 | 23.7 | 18.51 | 20.19 | — |

**Figure 5.**
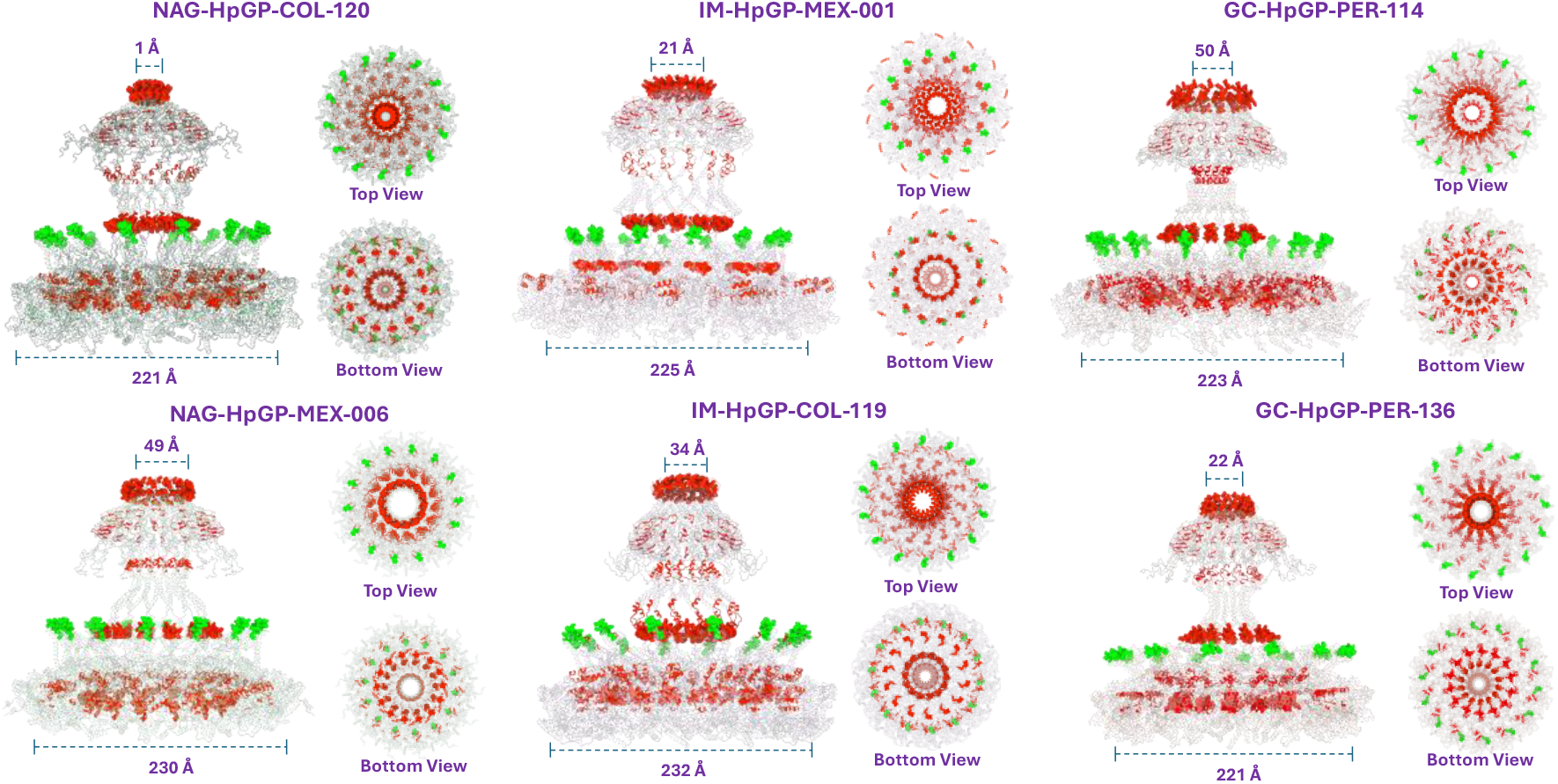
*HpGP* SWEurope CagY multimeric stabilized structures. The figure shows six structures of the CagY protein, where columns represent the phenotype and are color-coded accordingly. The GC-associated CagY structure displays a stretched pore. The IM-associated CagY structure shows the least reduction in pore size, while the NAG-associated CagY does not exhibit structural similarity.

Another approach was to model CagY from strains after monkey (rhesus macaque) passage of the J66 IL-8 inducer strain, as described previously by Solnick et al. (Barrozo et al., 2013). The output strain rOut1 did not induce the release of IL-8, whereas the rOut3 and J166 did induce IL-8, despite coming from the same IL-8 inducing strain and only after a brief monkey colonization. Of note, CagY from the non-IL-8 induction strain had marked differences in the virB10 sequence, particularly in the AP region that associated with a reduced dimension of the crown at the multimer tip and changes in the MRR region associated with marked structural differences at the base of the multimer (Figure 6). In both regions the changes led to a closed tunnel, which would cancel the translocation of molecules through the t4SS. In contrast, the IL8+ strain (J166-IL8+ and rOut3-IL8+) presented an opened tunnel, plus other differences in the rest of the multimer.

**Figure 6.**
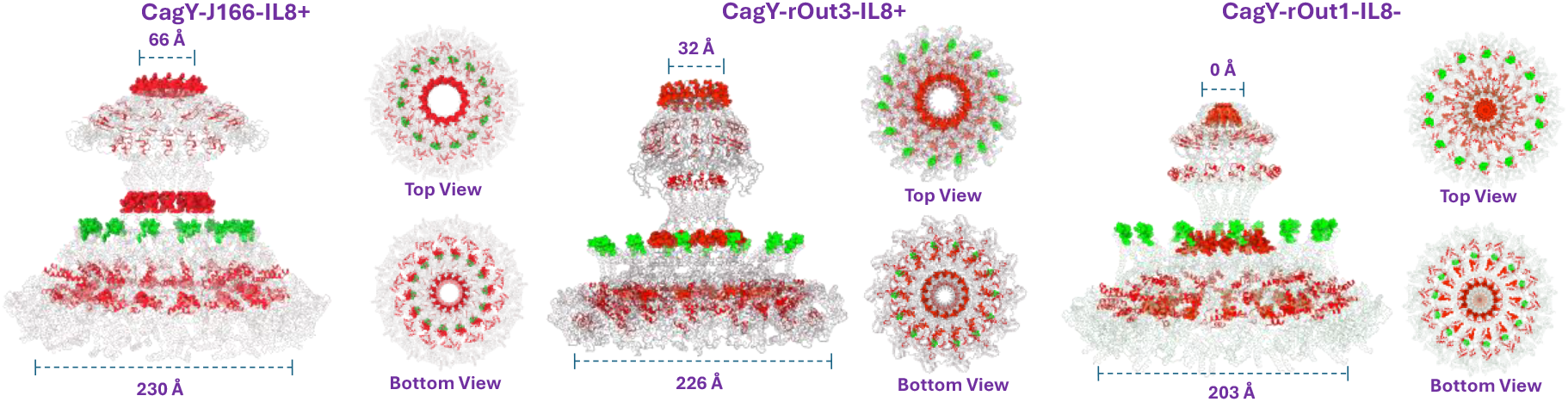
J166 CagY variants multimeric stabilized structures. The figure shows four structures of the CagY protein. The variants displays a high structure variability within structures, interestingly, the non-IL8 producer, shows pore channel fully closed, meanwhile, the IL8 producer CagY displays a fully open pore channel.

### Proline, a key amino acid for the structure of complex proteins is abundant in CagY

A highly specialized structure like CagY multimer that interact with DNA, proteins and even other metabolites, all with specialized functions, requires a very tight control of the structure and besides cysteine we wondered if the protein was rich in proline. This was the case and CagY presents an average of 50 prolines, most present in the MRR and in the VirB10 homolog regions, shaping rings in the MRR, AP and VirB10 regions (Figure 7). In contrast to cysteine present only in MRR, proline is also important in the structure of the VirB10 region, including AP. One proline is present in each of the A and B repeats and of note, like the cysteines, proline seems to be highly conserved.

**Figure 7.**
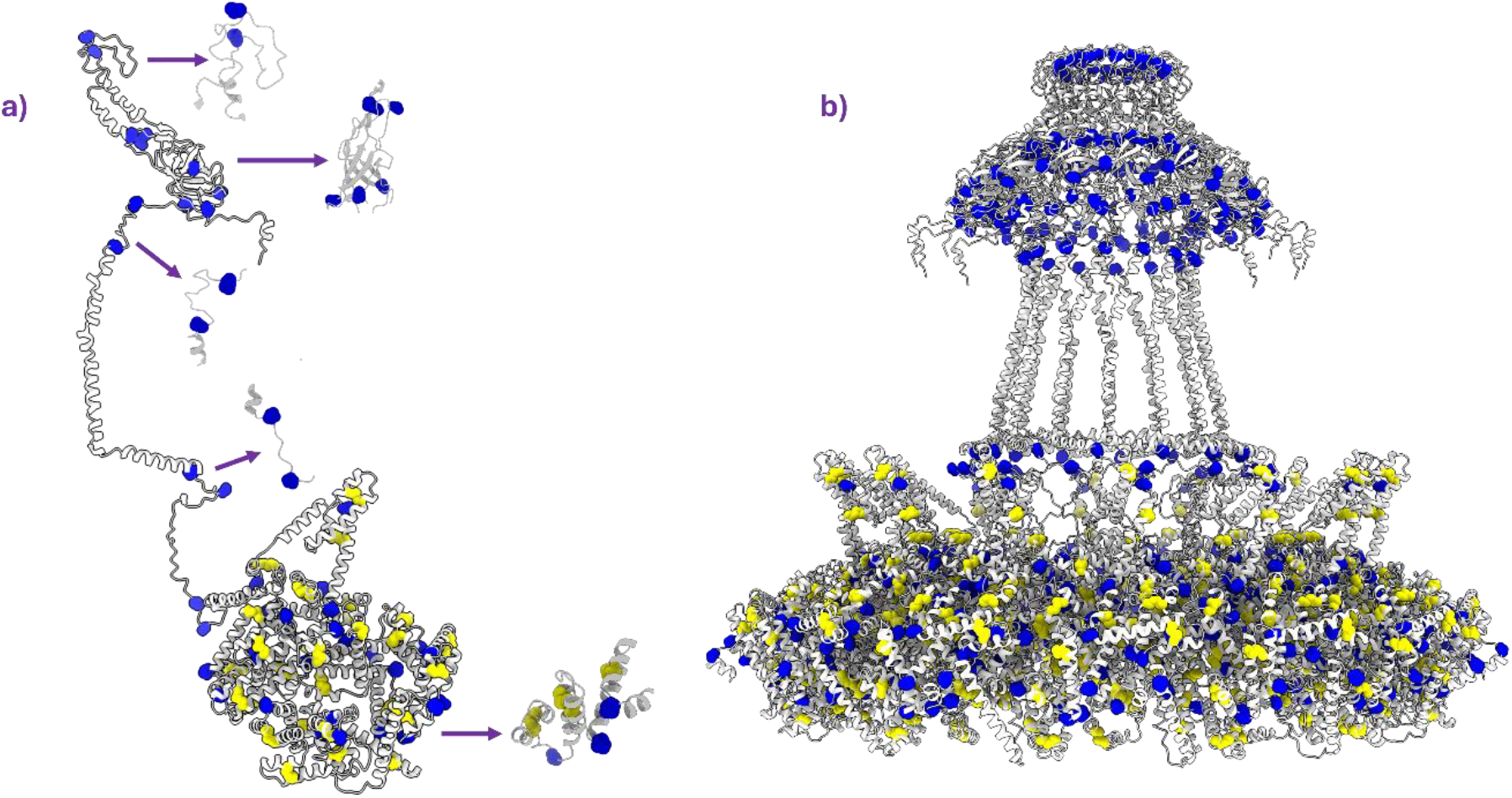
Three-dimensional representation of Cysteine and Proline aminoacid within CagY of *Hp* 26695. a) CagY monomer structure shows the Cysteine and Proline, this last is responsible of globular and VirB10 homologous region conformation as it’s shown in the arrowed regions. b) CagY multimer structure shown the high presence of Cysteine and Proline aminoacid.

Finally, we illustrated the cysteines and prolines in the same six sequences from patients (Figure 8). In the only strain showing closed the upper tip (NAG-HpGP-COL-120), proline in AP seems to be fully closing the tunnel. In the two GC strains showing stretched the neck an extra proline is in the more stretched spot. The same extra copy was present in an IM and a NAG strain but here the neck was not stretched. Major differences were observed in the distribution of proline in the MRR and VirB10 regions, associated with important structural changes, but no evident association with disease.

**Figure 8.**
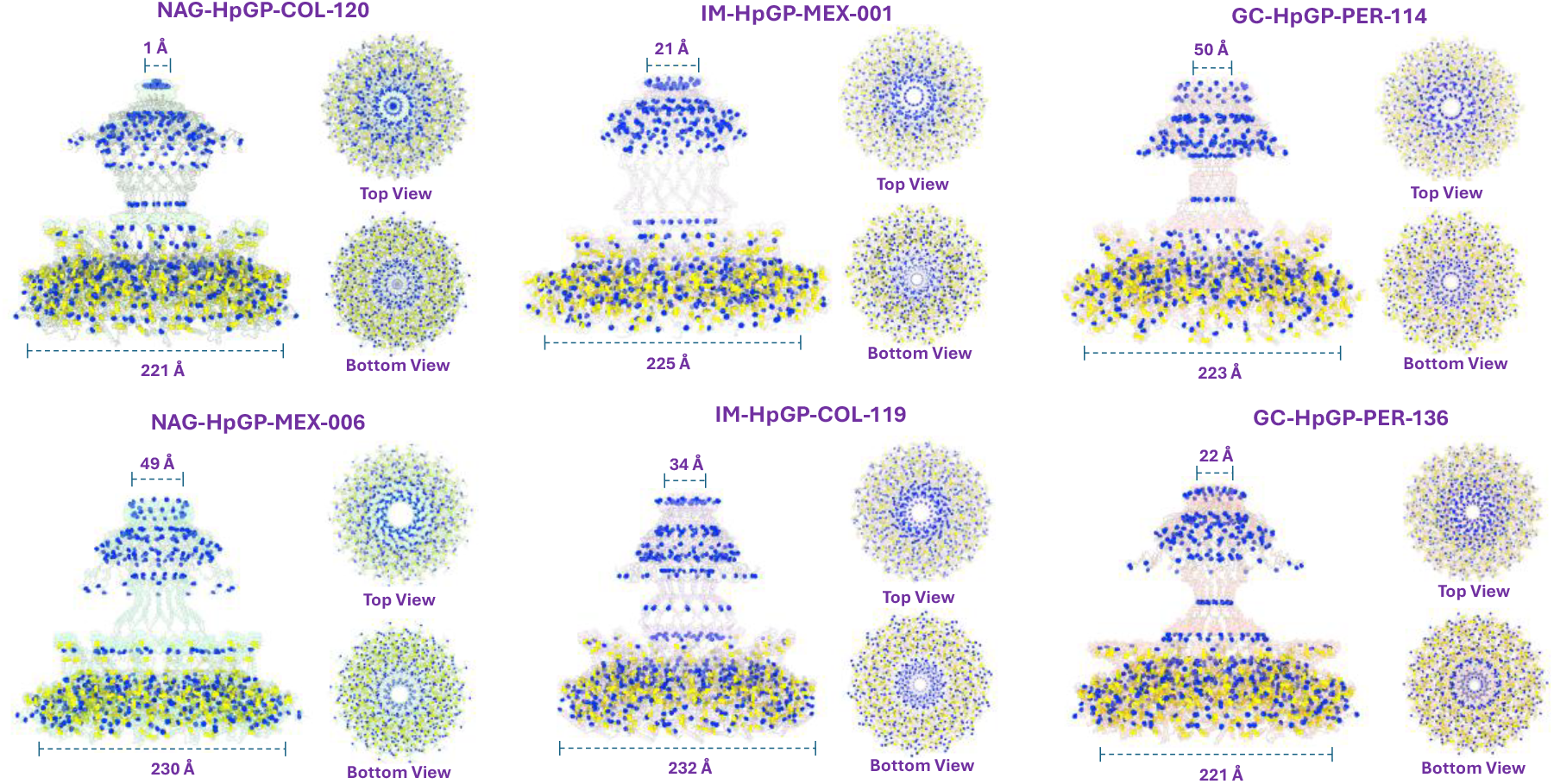
*HpGP* SWEurope CagY multimeric stabilized structures. The figure shows six structures of the CagY protein, where columns represent the phenotype and are color-coded accordingly. The structures displays the presence of cysteine and proline aminoacid.

In the strains before and after monkey passage of J166 (Figure 9) the prolines in the VirB10 and in the MRR regions varied significatively, formed a ring large enough to possible help maintain open the tunnel. In contrast, proline rings in the rOut1 IL8(-) strain had a very tight ring, favoring a closed tunnel (Figure 9).

**Figure 9.**
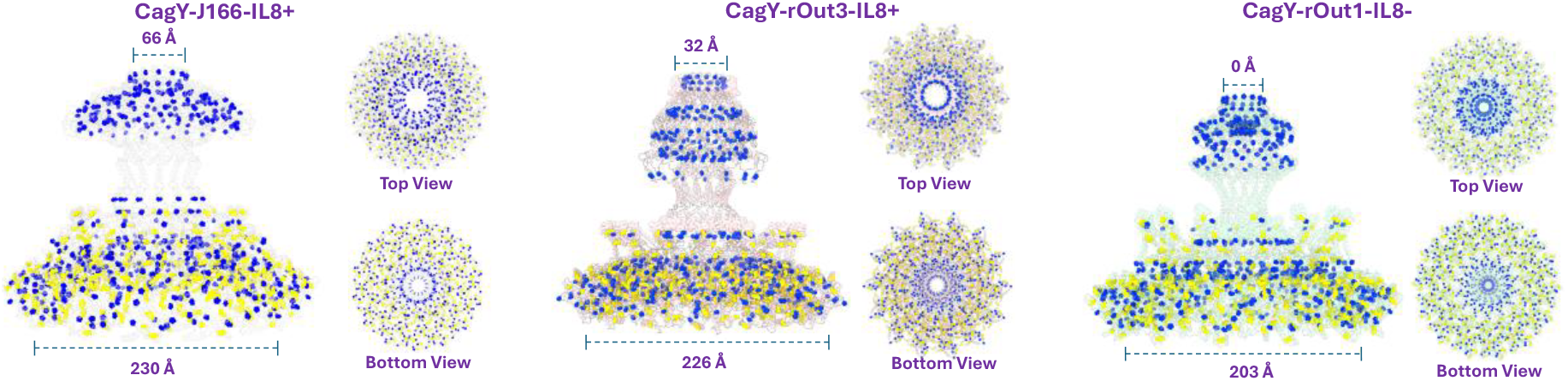
J166 CagY variants multimeric stabilized structures. The figure shows four structures of the CagY protein. The models display presence of cysteine and proline aminoacids

## Discussion

CagY has shown to be a highly complex protein, the largest protein in *Hp* and among the largest bacterial virulence proteins, with close to 2,000 aa, 800 of which are a large region of repeating units. It is a protein that traverses from the bacterial cytoplasm to the outside of the bacterium. Furthermore, the sequence of the gene is hard to assemble using short-reads sequencing because of the large number of repeat-sequences, and there is no known homologous gene among all described proteins (Delahay et al., 2008; López-Luis et al., 2023). Working with a large collection of *cagY* genes sequenced using long reads we were able to overcome previous troubles after short reads sequencing. Phylogeny, PCA and networking analyses confirmed the strong population structure of *cagY*, like what has been observed with the core genome (Yahara et al., 2013). We previously reported this high population structure in other virulence genes like *cagA* and *vacA (*Muñoz-Ramirez et al., *2021)*, which suggests these genes have coevolved with its human host, probably because they are under strong selective pressure due to its interaction with human proteins. Furthermore, the analyses limited to the MRR region also showed a clear population structure, indicating this is a CagY region under strong selective pressure to coevolve with its human host. It has been suggested that MRR is an important region to regulate the proinflammatory activity of the t4SS (Barrozo et al., 2016; Bella et al., 2021; Sierra et al., 2019) but its complexity has hindered proper functional analyses. We recently showed the importance of MRR in the structure of CagY because of its unusual high number of cysteines concentrated in this region of the protein (López-Luis et al., 2023), a density of S-S bridges not observed in any other protein that we are aware of, not even in mammals. High cysteine content has been found in high life domains as vertebrates, reptiles and plants, particularly in kinases or important for stress response and selective pressure, homeostasis and in immune response as well as in protein-structure (Barrozo et al., 2016; Delahay et al., 2008; Tran et al., 2023; Wang et al., 2014; Zhang et al., 2017). It is important to highlight that even in higher life domains 48 Cys in one protein is an extremely high and unusual number, making CagY an extraordinarily rare protein, with no known homology to any other protein (Barrozo et al., 2016; Pace & Weerapana, 2013). We recently suggested that these Cys residues might be important in the highly folded protein structure formed by the MRR region, but also for the structure of the multimer where 14 MRRs form the base of the donut and the tunnel for the translocation of molecules from HP to the epithelial cell (López-Luis et al., 2023). This means that as much as 672 (14X48) Cys residues are necessary to make this complex structure possible. The study of the origin and evolution of CagY is a challenging and exciting subject, even more if we consider that the HP T4SS evolved to interact specifically with human gastric epithelial cells and is the result of at least 100,000 years of co-evolution human-Hp (Falush et al., 2003). In this sense it is not a surprise the strong population structure of *cagY* and even of the MRR region of the gene.

It was important to understand the extent of population structure in the gene because when present, it may mask or confound any other analyses. Once we learned that common analytical methods reveal a strong population structure, we tried different approaches to analyze if there was a way to distinguish the clinical source of *cagY* by studying its sequence diversity. In short, the pipeline that worked was the following: a library with unitigs generated after fragmenting each *cagY* sequence was constructed and used to scan each of the 687 *cagY* genes studied followed by dimensional reduction analyses. Next, the unitigs that resulted informative to differentiate the clinical groups were selected with both, RandomForest (RF) and LDA. The results showed a clear separation of the genes according to disease, and although some of the selected unitigs were also informative for ancestry, many of them were useful to distinguish between GC, IM or NAG. Thus, we found that the *cagY* gene sequence does have information that may identify *Hp* strains with an increased risk of association with GC, a task that has been elusive in previous works. We built a supervised RF model using disease associated CagY fragments and assembled a variant database comprising those informative unitigs linked to clinical outcomes. It is important to highlight that almost 40% of the informative unitigs were also associated to ancestry, documenting the strong population structure of *cagY*.

Still, the other 60% unitigs were not associated with population structure and most of these included unitigs with the highest informative value.

With the above information, we were able to locate the position of each unitig associated with GC in the *cagY* sequence. It is interesting to note that most of the GC informative unitigs mapped to the MRR region, from the A8 to the A21 repeating-module and 25 mapped specifically in motifs δμ. Fifteen of the 25 included cysteine codons, indicating that they should be very relevant to the structure in both, the CagY monomer and the multimer and hence in the activity of the T4SS(Barrozo et al., 2016; Bella et al., 2021; Kylarova et al., 2016; Sierra et al., 2019). If the unusually high number of cysteines in the MRR region play a role other than in structure is something that needs to be further investigated (Zhang et al., 2017). It has also been suggested that the A module is responsible for the binding of CagY to α5β1 integrins, and recombination within this module may alter the binding (Skoog et al., 2018); however, we now know that the MRR is located in the periplasmic region and not exposed on the surface. This would then suggest that recombination within the A module may have an indirect effect on the binding to α5β1 integrins, probably as a result of modification of the protein structure.

Nine GC-associated unitigs located in the VirB10 homologous region, a region that includes the outer transmembrane region, but also sites of interaction with the other cagPAI proteins CagX and CagT (Skoog et al., 2018). Furthermore, VirB10 also includes the antenna region (AP) a fragment that is very important for the translocation of CagA to gastric epithelial cells, and hence in the regulation of IL8 induction (Tran et al., 2023). Thus, this small region of the CagY protein has multiple functions that might be sensitive to mutations and our results show variants that affect the risk for GC. In addition, the AP showed a strong and significant co-presence with other informative unitigs present in the MRR and FRR regions, suggesting that a modification in one region may affect other functional regions of this large multifunctional protein.

Considering the complex multidomain, multifunction and high sequence diversity nature of CagY (Delahay et al., 2008), searching for any structure-function relationship is a very challenging task. Yet, it is a task we should start pursuing and for that, we modelled CagY from NAG, IM and GC strains all with the same hspSWEuropeMiscAmerica ancestry to minimize the influence of the population. As expected, major differences between disease strains were observed along the 14-multimer, with difference in diameter of the central tunnel, in the structure of the MRR formed donut, or in the AP region at the tip of the multimer where interaction with cell receptors occurs (Tran et al., 2023). However, no evident disease-associated differences were observed, most probably because of the low number of sequences tested (two of each disease group) and so, a larger number of cases need to be modelled to find if there is any disease-associated pattern.

Another approach was to study CagY from the J166 *Hp* strain (IL-8 inducer) after inoculated into rhesus monkeys (Barrozo et al., 2013), where we modelled one output strain inducing IL8 and one that did not. A noticeable difference was that in the non-inducing strain the multimer presented the central tunnel closed (like one NAG did), whereas in the inducing strain the tunnel was open (Like in the two GC strains). Thus, changes in the lumen of the tunnel might be one mechanism to regulate the translocation of CagA and hence the induction of inflammation. Although other mechanisms, such as the alteration of the interaction of CagY with the other proteins of the T4SS might also be involved. Also, the cysteine residues could play a key regulatory function in CagY protein, thus, in our previous work (López-Luis et al., 2023), we discussed about how these residues contain reductor power, flexibility and could help to CagA translocation.

Prolines are known for their functional roles in transmembrane proteins such as transporters or ion channels (Sansom, 1992). Because of its properties, proline play an important role in the structure of membrane proteins facilitating formation and stabilization of helixes (Woolfson & Williams, 1990) and modeling studies suggest they may also function as molecular switches or hinges by allowing conformational changes (S.P. Sansom & Weinstein, 2000). These observations encouraged us to search for the presence of proline rich regions in CagY, since it is a transmembrane multimer that function as a channel for diverse molecules and has regions with densely packed helices arranged in concentering rings that may work as switch. Not surprisingly, we found these proline rich regions along the protein that in the multimer resulted in rings surrounding the channel at specific locations. It was more abundant in the MRR region regularly spaced between the cysteines, suggesting both amino acids are important in the structure and function of the CagY complex. The VirB10 homolog region was also proline enriched, including the apical AP site. As mentioned before, the modelled 14 CagY-multimer contains approximately 672 cysteine residues and may also include around 700 proline residues, an extraordinarily high concentration of these amino acids in a transmembrane transport system. This work is the first reporting a detail modelling of most of the backbone of the *HP* T4SS, a bacterial system highly specialized to work with human gastric epithelial cells.

In summary, we found that *cagY* gene present a strong population structure, which attest for an ancient co-evolution of the protein with humans, a fact that interferes with other analyses. Still, a proper informatic analyses of *cagY* variants may distinguish the clinical groups of NAG, IM and GC and identified the sequences of the short probes (unitigs) that distinguished these groups. The position of each probe in the protein was localized and found that most of the informative unitigs mapped in the complex repetitive region (MRR) of the gene where cysteine bridges and prolines are enriched. The modelling of some of the variants illustrated major changes in the structure of the 14-multimer complex that might be part of the mechanisms by which CagY regulate the function of the T4SS and hence modulate the inflammatory response.

## Acknowledgment

We thank Dr. Santiago Sandoval for his help to perform the phylogenetic network analyses of *cagY*.

## Supplementary Figures

**Supplementary Figure 1.**
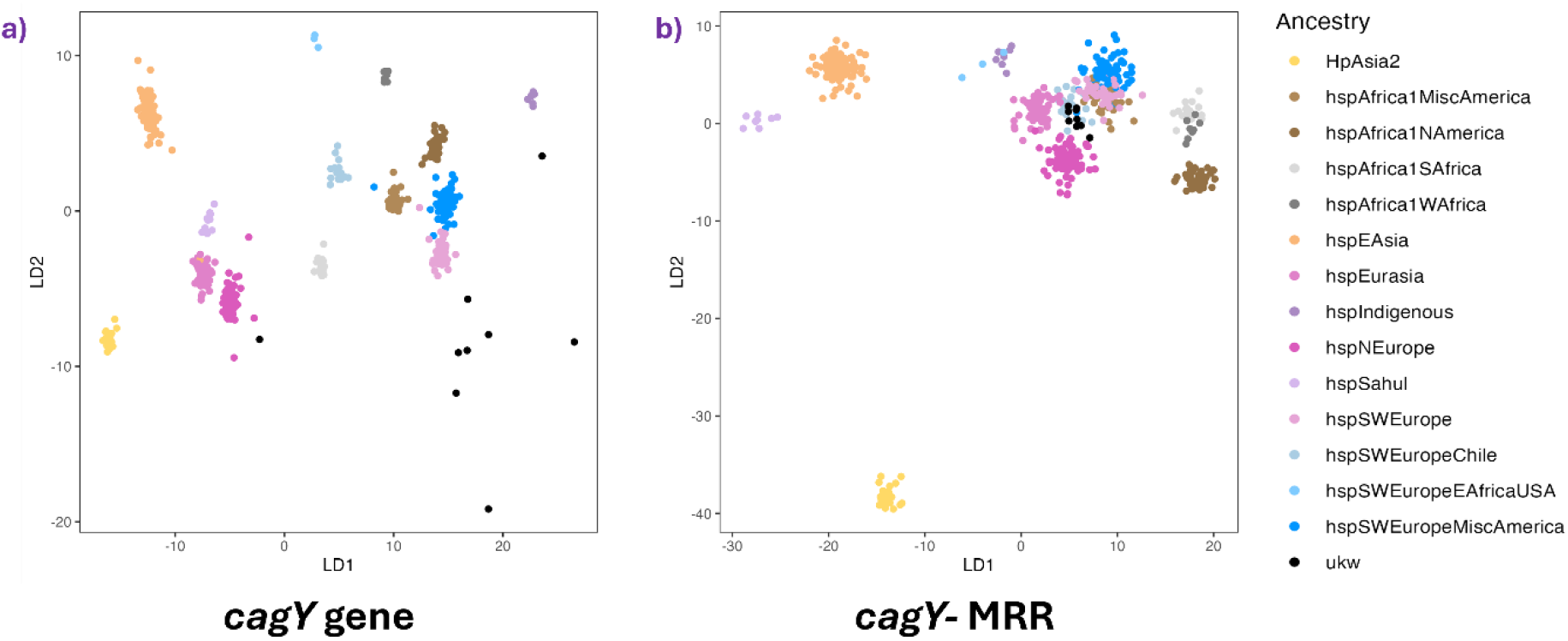
Linear Discriminant Analysis (LDA) of the *cagY* gene based on virtual hybridization. **a)** LDA of the complete *cagY* gene, where clusters are clearly defined. **b)** LDA of the *cagY*MRR region, showing lower resolution and the clustering of European-American sequences.

**Supplementary Figure 2.**
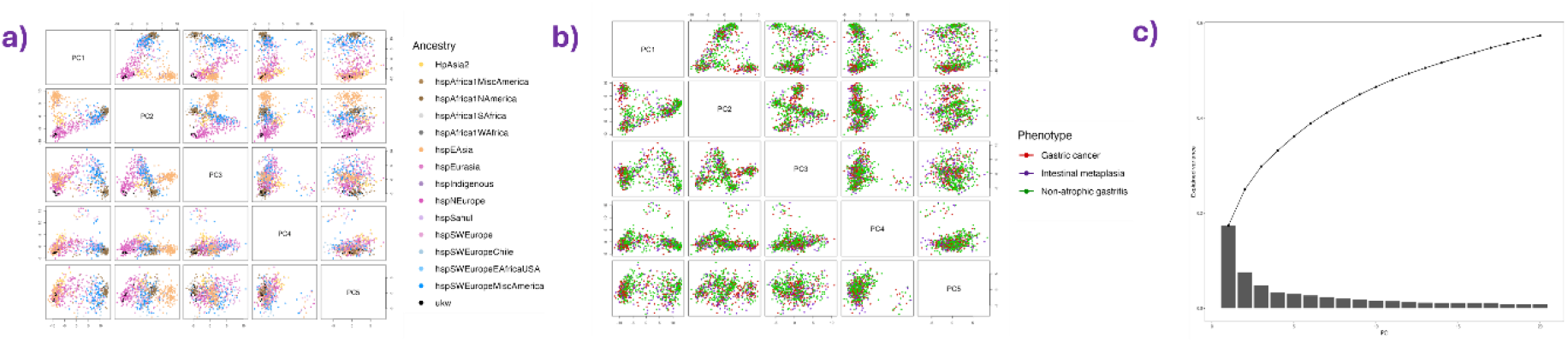
Principal Component Analysis (PCA) of the unitig presence/absence matrix. **a)** PCA of the first 5 principal components (PCs). The first 2 components show a clear association with the *H. pylori* population structure, while the last 3 components do not exhibit a relationship with the population structure of *H. pylori*. **b)** Explained variance of the PCA, showing the variance captured by the first 20 PCs.

**Supplementary Figure 3.**
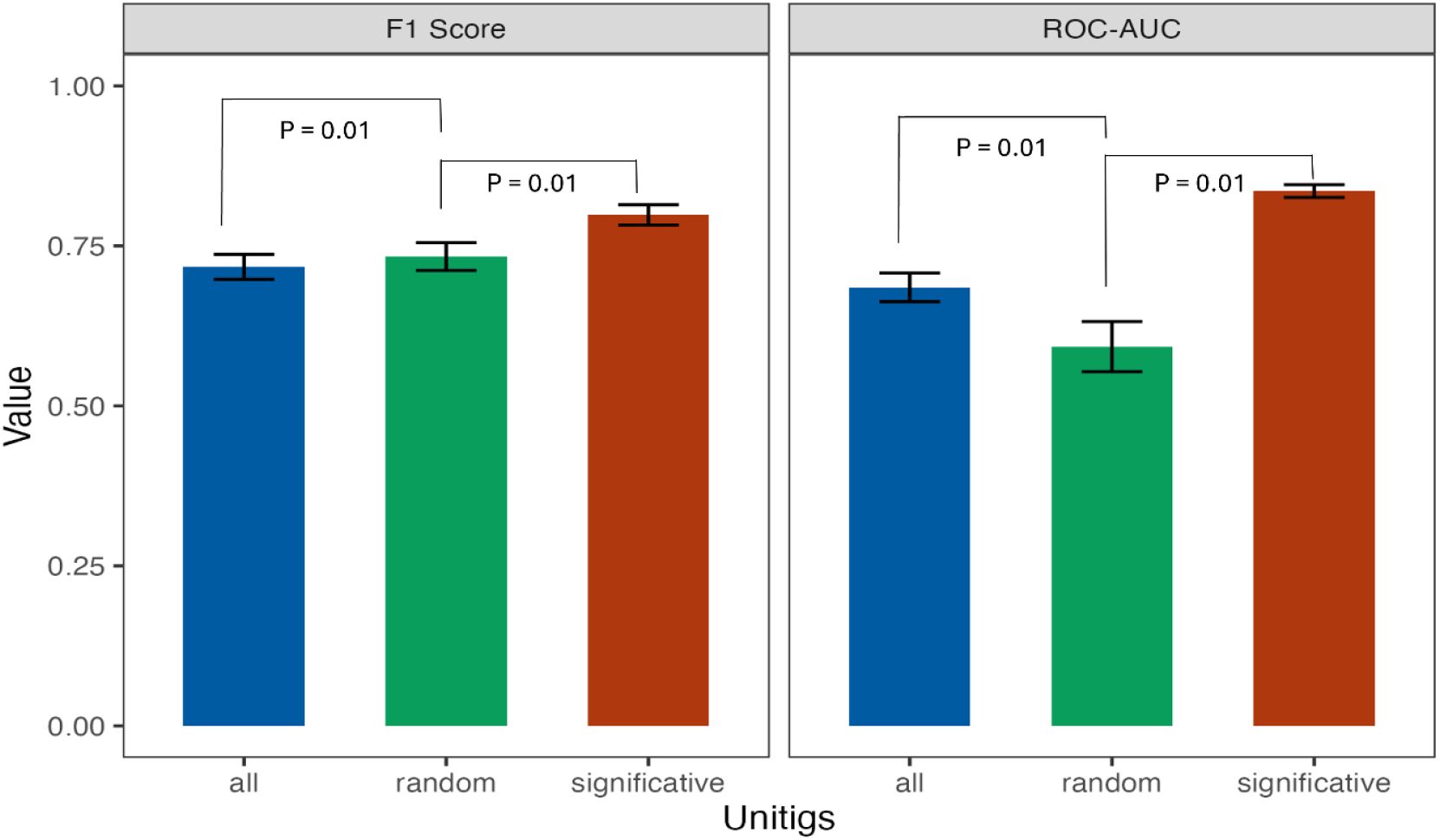
Comparison of three datasets of unitigs tested with the Random Forest algorithm. The figure shows two metrics obtained from training the three datasets: the F1 score and ROC-AUC. In both metrics, the best-performing models were constructed using the significant GC-associated unitigs.

**Supplementary Figure 4.**
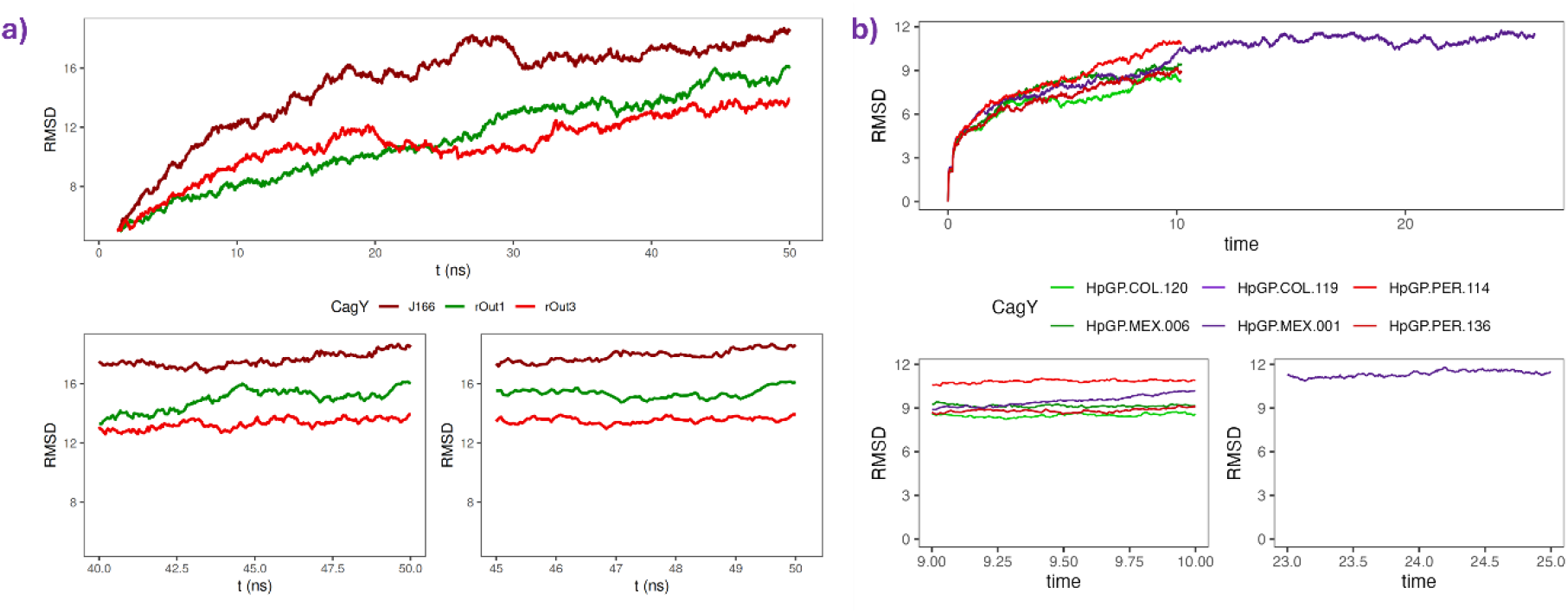
Root Mean Square Deviation (RMSD) plot of Molecular Dynamics (MD) stabilization. **a)** RMSD plot of 50 ns for the CagY models from Barrozo et al., 2013. **b)** RMSD plot of 10-20 ns for the *HpGP* SWEurope CagY models. The bottom plots show the stabilization during the final nanoseconds of the MD simulations.

**Supplementary Figure 5.**
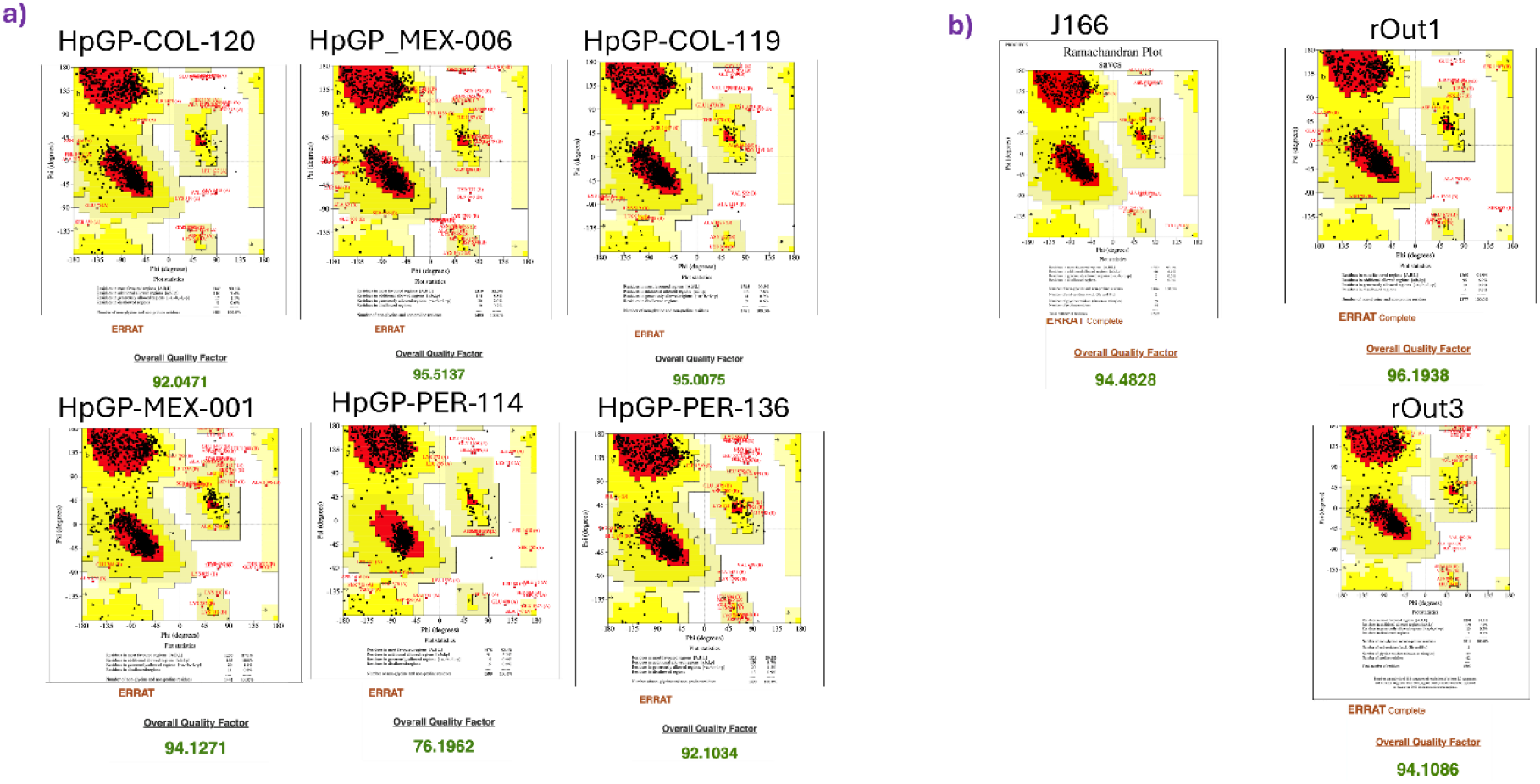
Validation analysis with Ramachandran plot (PROCHECK) and ERRAT server. **a)** Ramachandran plots of the CagY models from Barrozo et al., 2013. **b)** Ramachandran plots of the *HpGP* SWEurope CagY models.

**Supplementary Figure 6.**
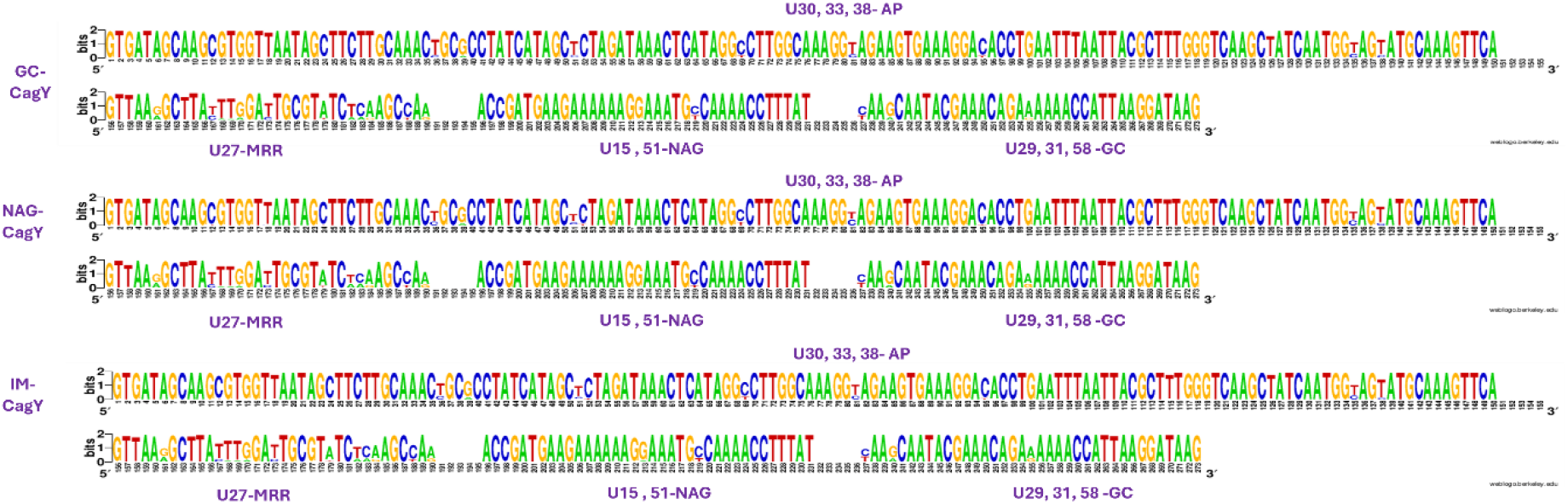
Nucleotide alignment of AP and the three most significative regions of *cagY* gene based *Hp* phenotype. The phenotype-based alignments show differences in the proportion of multiple nucleotides in the AP and GC and NAG which is associated with the differences of presence/absence. The MRR unitig (U27) display many variations among sequence.

**Supplementary Figure 7.**
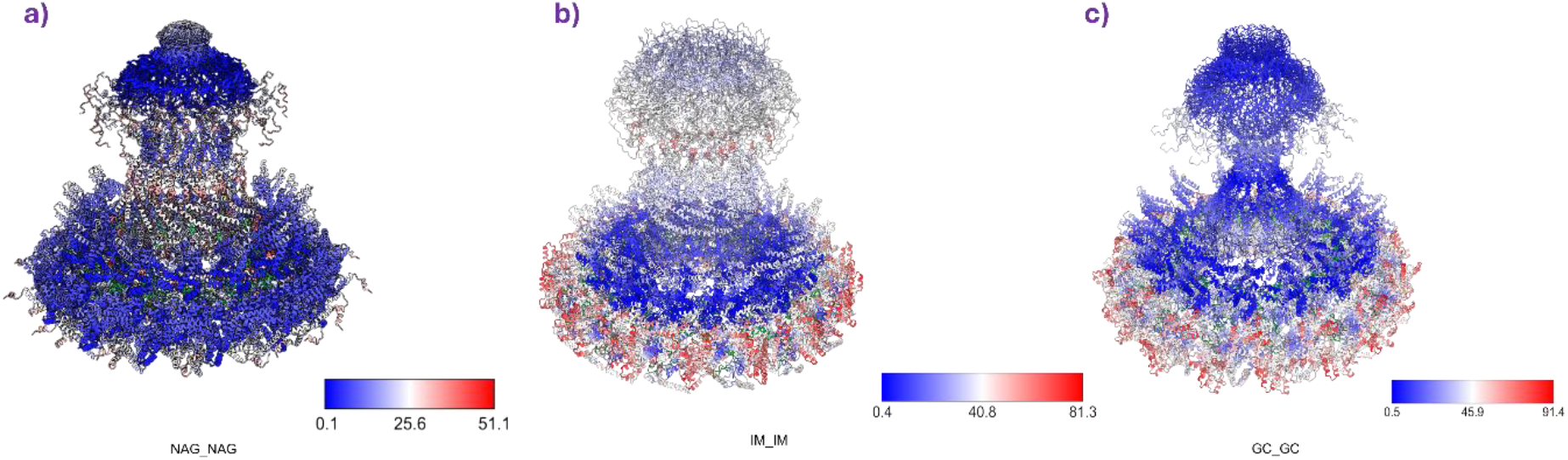
Root Mean Square Deviation comparison of CagY structures of same phenotype. NAG structures show structural homology in almost all protein, only shows differences in the connection of VirB10 homology region with MRR. IM structures show different in most of the structure, only in the inner pore of MRR the structures are similar. The GC structures show structural differences in the MRR on the outside of CagY.

## Supplementary Tables

**Supplementary Table 1.** CagY gene status of the HpGP set

| **HpGP ID** | **Country** | **cagY status**<br>**by NCBI**<br>**annotation** | **GenBank**<br>**Accession ID** |
| --- | --- | --- | --- |
| **HpGP-26695-**<br>**ATCC** | UK | + | CP079087 |
| **HpGP-ALG-001** | Algeria | - | CP079086 |
| **HpGP-ALG-002** | Algeria | - | CP079085 |
| **HpGP-ALG-003** | Algeria | + | CP079084 |
| **HpGP-ALG-004** | Algeria | - | CP079083 |
| **HpGP-ALG-005** | Algeria | - | CP079093 |
| **HpGP-ALG-006** | Algeria | - | CP079082 |
| **HpGP-ALG-007** | Algeria | - | CP079081 |
| **HpGP-ALG-008** | Algeria | + | CP079080 |
| **HpGP-ALG-009** | Algeria | - | CP079079 |
| **HpGP-ALG-010** | Algeria | + | CP079078 |
| **HpGP-ARG-001** | Argentina | + | CP079077 |
| **HpGP-ARG-002** | Argentina | - | CP080089 |
| **HpGP-ARG-003** | Argentina | - | CP079076 |
| **HpGP-ARG-004** | Argentina | - | CP079075 |
| **HpGP-ARG-005** | Argentina | - | CP079074 |
| **HpGP-ARG-006** | Argentina | + | CP079073 |
| **HpGP-ARG-007** | Argentina | + | CP079072 |
| **HpGP-ARG-008** | Argentina | + | CP079071 |
| **HpGP-ARG-009** | Argentina | + | CP079070 |
| **HpGP-ARG-010** | Argentina | + | CP079069 |
| **HpGP-BGD-001** | Bangladesh | + | CP079068 |
| **HpGP-BGD-002** | Bangladesh | + | CP079067 |
| **HpGP-BGD-003** | Bangladesh | + | CP079092 |
| **HpGP-BGD-004** | Bangladesh | + | CP079066 |
| **HpGP-BGD-005** | Bangladesh | + | CP079091 |
| **HpGP-BGD-006** | Bangladesh | - | CP079065 |
| **HpGP-BGD-007** | Bangladesh | - | CP079064 |
| **HpGP-BGD-008** | Bangladesh | + | CP079063 |
| **HpGP-BGD-009** | Bangladesh | + | CP079062 |
| **HpGP-BGD-010** | Bangladesh | - | CP079061 |
| **HpGP-BGR-001** | Bulgaria | + | CP079060 |
| **HpGP-BGR-003** | Bulgaria | + | CP079059 |
| **HpGP-BGR-004** | Bulgaria | - | CP079058 |
| **HpGP-BGR-005** | Bulgaria | + | CP079057 |
| **HpGP-BGR-006** | Bulgaria | + | CP079056 |
| **HpGP-BGR-007** | Bulgaria | + | CP079055 |
| **HpGP-BGR-008** | Bulgaria | - | CP079054 |
| **HpGP-BGR-010** | Bulgaria | - | CP079053 |
| **HpGP-BRA-001** | Brazil | + | CP079052 |
| **HpGP-BRA-002** | Brazil | + | CP079051 |
| **HpGP-BRA-003** | Brazil | - | CP079050 |
| **HpGP-BRA-004** | Brazil | + | CP079049 |
| **HpGP-BRA-005** | Brazil | + | CP079048 |
| **HpGP-BRA-006** | Brazil | - | CP079047 |
| **HpGP-BRA-007** | Brazil | + | CP079046 |
| **HpGP-BRA-008** | Brazil | + | CP079045 |
| **HpGP-BRA-009** | Brazil | - | CP079044 |
| **HpGP-BRA-010** | Brazil | + | CP079043 |
| **HpGP-BRA-012** | Brazil | + | CP079042 |
| **HpGP-BRA-013** | Brazil | + | CP079041 |
| **HpGP-BRA-014** | Brazil | + | CP079040 |
| **HpGP-BRA-015** | Brazil | + | CP079039 |
| **HpGP-BRA-017** | Brazil | - | CP079038 |
| **HpGP-BRA-018** | Brazil | + | CP079037 |
| **HpGP-BRA-019** | Brazil | + | CP079036 |
| **HpGP-BRA-020** | Brazil | + | CP079035 |
| **HpGP-BRA-022** | Brazil | + | CP079034 |
| **HpGP-BRA-023** | Brazil | + | CP079033 |
| **HpGP-BRA-025** | Brazil | + | CP079032 |
| **HpGP-CAN-001** | Canada | - | CP079031 |
| **HpGP-CAN-002** | Canada | + | CP079090 |
| **HpGP-CAN-004** | Canada | + | CP079030 |
| **HpGP-CAN-006** | Canada | - | CP079029 |
| **HpGP-CAN-007** | Canada | - | CP079028 |
| **HpGP-CAN-008** | Canada | + | CP079027 |
| **HpGP-CAN-010** | Canada | - | CP079026 |
| **HpGP-CAN-011** | Canada | - | CP079025 |
| **HpGP-CAN-012** | Canada | + | CP079024 |
| **HpGP-CAN-014** | Canada | + | CP079023 |
| **HpGP-CAN-015** | Canada | - | CP079022 |
| **HpGP-CAN-016** | Canada | - | CP079021 |
| **HpGP-CAN-017** | Canada | + | CP079020 |
| **HpGP-CAN-018** | Canada | - | CP079019 |
| **HpGP-CAN-019** | Canada | - | CP079018 |
| **HpGP-CAN-020** | Canada | - | CP079017 |
| **HpGP-CAN-022** | Canada | + | CP079016 |
| **HpGP-CAN-023** | Canada | - | CP079015 |
| **HpGP-CAN-025** | Canada | + | CP079014 |
| **HpGP-CAN-026** | Canada | + | CP079013 |
| **HpGP-CHI-001** | Chile | - | CP079598 |
| **HpGP-CHI-002** | Chile | - | CP079597 |
| **HpGP-CHI-004** | Chile | + | CP079596 |
| **HpGP-CHI-005** | Chile | + | CP079595 |
| **HpGP-CHI-006** | Chile | - | CP079594 |
| **HpGP-CHI-007** | Chile | - | CP079593 |
| **HpGP-CHI-008** | Chile | - | CP079592 |
| **HpGP-CHI-010** | Chile | - | CP082183 |
| **HpGP-CHI-011** | Chile | + | CP079591 |
| **HpGP-CHI-014** | Chile | - | CP079590 |
| **HpGP-CHI-016** | Chile | - | CP079589 |
| **HpGP-CHI-017** | Chile | - | CP079588 |
| **HpGP-CHI-018** | Chile | + | CP079587 |
| **HpGP-CHI-021** | Chile | + | CP079586 |
| **HpGP-CHI-022** | Chile | + | CP079012 |
| **HpGP-CHI-023** | Chile | + | CP079585 |
| **HpGP-CHI-024** | Chile | + | CP079584 |
| **HpGP-CHI-026** | Chile | + | CP079583 |
| **HpGP-CHI-027** | Chile | + | CP079582 |
| **HpGP-CHI-028** | Chile | + | CP079581 |
| **HpGP-CHI-101** | Chile | - | CP079011 |
| **HpGP-CHI-102** | Chile | - | CP079580 |
| **HpGP-CHI-103** | Chile | - | CP079010 |
| **HpGP-CHI-104** | Chile | - | CP079579 |
| **HpGP-CHI-105** | Chile | - | CP079578 |
| **HpGP-CHI-106** | Chile | - | CP079577 |
| **HpGP-CHI-107** | Chile | - | CP079576 |
| **HpGP-CHI-110** | Chile | - | CP082182 |
| **HpGP-CHI-113** | Chile | - | CP079575 |
| **HpGP-CHI-114** | Chile | - | CP079574 |
| **HpGP-CHI-116** | Chile | - | CP079573 |
| **HpGP-CHI-117** | Chile | - | CP079572 |
| **HpGP-CHI-119** | Chile | + | CP079571 |
| **HpGP-CHI-120** | Chile | + | CP079570 |
| **HpGP-CHI-122** | Chile | + | CP079569 |
| **HpGP-CHI-123** | Chile | + | CP079568 |
| **HpGP-CHI-124** | Chile | + | CP079567 |
| **HpGP-CHI-125** | Chile | + | CP079566 |
| **HpGP-CHI-126** | Chile | - | CP079565 |
| **HpGP-CHI-127** | Chile | + | CP079564 |
| **HpGP-CHI-204** | Chile | - | CP079009 |
| **HpGP-CHI-207** | Chile | - | CP079008 |
| **HpGP-CHI-208** | Chile | + | CP079007 |
| **HpGP-CHI-209** | Chile | + | CP079006 |
| **HpGP-CHI-210** | Chile | - | CP079005 |
| **HpGP-CHI-212** | Chile | + | CP079004 |
| **HpGP-CHN-001** | China | + | CP079003 |
| **HpGP-CHN-002** | China | + | CP079002 |
| **HpGP-CHN-003** | China | + | CP079001 |
| **HpGP-CHN-004** | China | + | CP079000 |
| **HpGP-CHN-005** | China | + | CP078999 |
| **HpGP-CHN-006** | China | + | CP078998 |
| **HpGP-CHN-007** | China | + | CP078997 |
| **HpGP-CHN-008** | China | + | CP078996 |
| **HpGP-CHN-009** | China | + | CP078995 |
| **HpGP-CHN-010** | China | + | CP078994 |
| **HpGP-COG-001** | DR Congo | + | CP078993 |
| **HpGP-COG-002** | DR Congo | + | CP078992 |
| **HpGP-COG-003** | DR Congo | + | CP078991 |
| **HpGP-COG-004** | DR Congo | + | CP078990 |
| **HpGP-COG-005** | DR Congo | + | CP078989 |
| **HpGP-COG-006** | DR Congo | + | CP078988 |
| **HpGP-COG-007** | DR Congo | + | CP078987 |
| **HpGP-COG-008** | DR Congo | + | CP078986 |
| **HpGP-COG-009** | DR Congo | + | CP078985 |
| **HpGP-COG-010** | DR Congo | + | CP078984 |
| **HpGP-COG-011** | DR Congo | + | CP078983 |
| **HpGP-COL-002** | Colombia | - | CP078982 |
| **HpGP-COL-003** | Colombia | + | CP078981 |
| **HpGP-COL-004** | Colombia | - | CP078980 |
| **HpGP-COL-005** | Colombia | - | CP078979 |
| **HpGP-COL-006** | Colombia | + | CP078978 |
| **HpGP-COL-008** | Colombia | + | CP078977 |
| **HpGP-COL-010** | Colombia | - | CP078976 |
| **HpGP-COL-102** | Colombia | + | CP078975 |
| **HpGP-COL-103** | Colombia | - | CP078974 |
| **HpGP-COL-104** | Colombia | + | CP078973 |
| **HpGP-COL-105** | Colombia | - | CP078972 |
| **HpGP-COL-106** | Colombia | - | CP078971 |
| **HpGP-COL-107** | Colombia | + | CP078970 |
| **HpGP-COL-108** | Colombia | + | CP078969 |
| **HpGP-COL-109** | Colombia | + | CP078968 |
| **HpGP-COL-110** | Colombia | + | CP078967 |
| **HpGP-COL-112** | Colombia | + | CP078966 |
| **HpGP-COL-113** | Colombia | + | CP078965 |
| **HpGP-COL-114** | Colombia | + | CP078964 |
| **HpGP-COL-118** | Colombia | + | CP078961 |
| **HpGP-COL-119** | Colombia | + | CP078960 |
| **HpGP-COL-120** | Colombia | + | CP078959 |
| **HpGP-COL-121** | Colombia | + | CP078958 |
| **HpGP-COL-122** | Colombia | + | CP078957 |
| **HpGP-COL-201** | Colombia | - | CP078956 |
| **HpGP-COL-202** | Colombia | - | CP078955 |
| **HpGP-COL-203** | Colombia | - | CP078954 |
| **HpGP-COL-204** | Colombia | + | CP078953 |
| **HpGP-COL-205** | Colombia | + | CP078952 |
| **HpGP-COL-206** | Colombia | + | CP078951 |
| **HpGP-COL-207** | Colombia | + | CP078950 |
| **HpGP-COL-208** | Colombia | + | CP078949 |
| **HpGP-COL-209** | Colombia | - | CP078948 |
| **HpGP-COL-210** | Colombia | - | CP078947 |
| **HpGP-COL-211** | Colombia | + | CP078946 |
| **HpGP-COL-301** | Colombia | + | CP078945 |
| **HpGP-COL-302** | Colombia | + | CP078944 |
| **HpGP-COL-303** | Colombia | - | CP078943 |
| **HpGP-COL-305** | Colombia | - | CP078942 |
| **HpGP-COL-306** | Colombia | + | CP078941 |
| **HpGP-COL-307** | Colombia | + | CP078940 |
| **HpGP-COL-308** | Colombia | + | CP078939 |
| **HpGP-COL-309** | Colombia | + | CP078938 |
| **HpGP-COL-310** | Colombia | - | CP078937 |
| **HpGP-COL-318** | Colombia | + | CP078936 |
| **HpGP-CRI-001** | Costa Rica | + | CP078935 |
| **HpGP-CRI-002** | Costa Rica | + | CP078934 |
| **HpGP-CRI-003** | Costa Rica | + | CP078933 |
| **HpGP-CRI-004** | Costa Rica | + | CP078932 |
| **HpGP-CRI-005** | Costa Rica | + | CP078931 |
| **HpGP-CRI-006** | Costa Rica | + | CP078930 |
| **HpGP-CRI-007** | Costa Rica | + | CP078929 |
| **HpGP-CRI-008** | Costa Rica | + | CP078928 |
| **HpGP-DOM-001** | Dominican Republic | - | CP078927 |
| **HpGP-DOM-002** | Dominican Republic | - | CP079089 |
| **HpGP-DOM-003** | Dominican Republic | - | CP078926 |
| **HpGP-DOM-004** | Dominican Republic | + | CP078925 |
| **HpGP-DOM-005** | Dominican Republic | + | CP078924 |
| **HpGP-DOM-006** | Dominican Republic | - | CP078923 |
| **HpGP-DOM-007** | Dominican Republic | - | CP078922 |
| **HpGP-DOM-008** | Dominican Republic | - | CP079088 |
| **HpGP-DOM-009** | Dominican Republic | - | CP078921 |
| **HpGP-DOM-010** | Dominican Republic | - | CP078920 |
| **HpGP-DOM-011** | Dominican Republic | + | CP078919 |
| **HpGP-FRA-001** | France | + | CP079563 |
| **HpGP-FRA-002** | France | + | CP079562 |
| **HpGP-FRA-003** | France | - | CP078918 |
| **HpGP-FRA-004** | France | - | CP082181 |
| **HpGP-FRA-005** | France | - | CP079561 |
| **HpGP-FRA-006** | France | - | CP079560 |
| **HpGP-FRA-007** | France | + | CP079559 |
| **HpGP-FRA-008** | France | - | CP079558 |
| **HpGP-FRA-009** | France | - | CP079557 |
| **HpGP-FRA-010** | France | - | CP079556 |
| **HpGP-FRA-011** | France | + | CP079555 |
| **HpGP-FRA-012** | France | + | CP079554 |
| **HpGP-FRA-013** | France | + | CP079553 |
| **HpGP-FRA-014** | France | + | CP079552 |
| **HpGP-FRA-015** | France | - | CP092856 |
| **HpGP-FRA-016** | France | + | CP079550 |
| **HpGP-FRA-017** | France | + | CP079549 |
| **HpGP-FRA-018** | France | + | CP078917 |
| **HpGP-FRA-019** | France | + | CP079548 |
| **HpGP-FRA-020** | France | + | CP079547 |
| **HpGP-FRA-021** | France | + | CP078916 |
| **HpGP-GER-001** | Germany | + | CP078915 |
| **HpGP-GER-002** | Germany | + | CP078914 |
| **HpGP-GER-003** | Germany | - | CP078913 |
| **HpGP-GER-004** | Germany | - | CP078912 |
| **HpGP-GER-005** | Germany | + | CP078911 |
| **HpGP-GER-006** | Germany | - | CP078910 |
| **HpGP-GER-007** | Germany | + | CP078909 |
| **HpGP-GER-009** | Germany | + | CP078908 |
| **HpGP-GER-010** | Germany | - | CP078907 |
| **HpGP-GER-012** | Germany | + | CP078906 |
| **HpGP-GER-013** | Germany | - | CP078905 |
| **HpGP-GER-015** | Germany | + | CP078904 |
| **HpGP-GER-016** | Germany | + | CP078903 |
| **HpGP-GER-018** | Germany | - | CP078902 |
| **HpGP-GER-019** | Germany | - | CP078901 |
| **HpGP-GER-020** | Germany | - | CP078900 |
| **HpGP-GER-021** | Germany | + | CP078899 |
| **HpGP-GHA-003** | Ghana | - | CP078898 |
| **HpGP-GHA-009** | Ghana | + | CP078897 |
| **HpGP-GMB-003** | The Gambia | - | CP078896 |
| **HpGP-GMB-004** | The Gambia | + | CP078895 |
| **HpGP-GMB-007** | The Gambia | + | CP078894 |
| **HpGP-GMB-008** | The Gambia | + | CP078893 |
| **HpGP-GMB-009** | The Gambia | + | CP078892 |
| **HpGP-GRE-001** | Greece | + | CP079546 |
| **HpGP-GRE-002** | Greece | - | CP079545 |
| **HpGP-GRE-003** | Greece | - | CP079544 |
| **HpGP-GRE-004** | Greece | - | CP079543 |
| **HpGP-GRE-005** | Greece | - | CP079542 |
| **HpGP-GRE-006** | Greece | - | CP079541 |
| **HpGP-GRE-008** | Greece | + | CP082180 |
| **HpGP-GRE-011** | Greece | - | CP079540 |
| **HpGP-GRE-012** | Greece | + | CP079539 |
| **HpGP-GRE-018** | Greece | + | CP078891 |
| **HpGP-GRE-019** | Greece | + | CP079538 |
| **HpGP-GRE-020** | Greece | - | CP079537 |
| **HpGP-GRE-022** | Greece | + | CP078890 |
| **HpGP-GRE-036** | Greece | - | CP079536 |
| **HpGP-GRE-039** | Greece | - | CP078889 |
| **HpGP-GRE-040** | Greece | + | CP078888 |
| **HpGP-GRE-041** | Greece | - | CP078887 |
| **HpGP-GRE-042** | Greece | - | CP078886 |
| **HpGP-GRE-043** | Greece | + | CP078885 |
| **HpGP-GRE-044** | Greece | + | CP078884 |
| **HpGP-GRE-046** | Greece | - | CP078883 |
| **HpGP-GTM-004** | Guatemala | + | CP082187 |
| **HpGP-GTM-007** | Guatemala | + | CP078882 |
| **HpGP-GTM-010** | Guatemala | + | CP078881 |
| **HpGP-HON-001** | Honduras | + | CP079535 |
| **HpGP-HON-002** | Honduras | + | CP079534 |
| **HpGP-HON-003** | Honduras | + | CP079533 |
| **HpGP-HON-004** | Honduras | + | CP079532 |
| **HpGP-HON-005** | Honduras | + | CP079531 |
| **HpGP-HON-006** | Honduras | + | CP079530 |
| **HpGP-HON-007** | Honduras | + | CP079529 |
| **HpGP-HON-009** | Honduras | + | CP079527 |
| **HpGP-HON-010** | Honduras | + | CP079526 |
| **HpGP-HON-011** | Honduras | + | CP079525 |
| **HpGP-HON-012** | Honduras | + | CP079524 |
| **HpGP-HON-013** | Honduras | + | CP079523 |
| **HpGP-HON-014** | Honduras | + | CP079522 |
| **HpGP-HON-015** | Honduras | + | CP079521 |
| **HpGP-HON-016** | Honduras | + | CP079520 |
| **HpGP-HON-017** | Honduras | + | CP079519 |
| **HpGP-HON-018** | Honduras | + | CP079518 |
| **HpGP-HON-019** | Honduras | + | CP079517 |
| **HpGP-HON-020** | Honduras | + | CP079516 |
| **HpGP-HON-021** | Honduras | + | CP079515 |
| **HpGP-HON-023** | Honduras | + | CP079513 |
| **HpGP-HON-024** | Honduras | - | CP079512 |
| **HpGP-HON-025** | Honduras | + | CP079511 |
| **HpGP-HON-026** | Honduras | + | CP079510 |
| **HpGP-HON-027** | Honduras | + | CP079509 |
| **HpGP-HON-028** | Honduras | + | CP079508 |
| **HpGP-IDN-001** | Indonesia | + | CP078880 |
| **HpGP-IDN-002** | Indonesia | + | CP078879 |
| **HpGP-IDN-003** | Indonesia | - | CP078878 |
| **HpGP-IDN-004** | Indonesia | + | CP078877 |
| **HpGP-IDN-005** | Indonesia | + | CP078876 |
| **HpGP-IDN-006** | Indonesia | - | CP078875 |
| **HpGP-IDN-007** | Indonesia | + | CP078874 |
| **HpGP-IDN-008** | Indonesia | + | CP078873 |
| **HpGP-IDN-009** | Indonesia | + | CP078872 |
| **HpGP-IDN-010** | Indonesia | - | CP078871 |
| **HpGP-IDN-011** | Indonesia | + | CP078870 |
| **HpGP-IND-001** | India | + | CP078869 |
| **HpGP-IND-002** | India | + | CP078868 |
| **HpGP-IND-003** | India | + | CP078867 |
| **HpGP-IND-004** | India | - | CP078866 |
| **HpGP-IND-005** | India | + | CP078865 |
| **HpGP-IND-006** | India | + | CP078864 |
| **HpGP-IND-007** | India | + | CP078863 |
| **HpGP-IND-008** | India | + | CP078862 |
| **HpGP-IND-009** | India | + | CP078861 |
| **HpGP-IND-010** | India | + | CP078860 |
| **HpGP-IRN-001** | Iran | + | CP078859 |
| **HpGP-IRN-002** | Iran | - | CP078858 |
| **HpGP-IRN-003** | Iran | - | CP078857 |
| **HpGP-IRN-009** | Iran | - | CP078856 |
| **HpGP-ISL-001** | Iceland | - | CP078855 |
| **HpGP-ISL-002** | Iceland | - | CP078854 |
| **HpGP-ISL-003** | Iceland | - | CP078853 |
| **HpGP-ISL-004** | Iceland | - | CP078852 |
| **HpGP-ISL-005** | Iceland | - | CP078851 |
| **HpGP-ISL-006** | Iceland | + | CP078850 |
| **HpGP-ISL-007** | Iceland | - | CP078849 |
| **HpGP-ISL-008** | Iceland | - | CP078848 |
| **HpGP-ISL-009** | Iceland | - | CP078847 |
| **HpGP-ISL-010** | Iceland | - | CP078846 |
| **HpGP-ISL-011** | Iceland | + | CP078845 |
| **HpGP-ISR-001** | Israel | - | CP078844 |
| **HpGP-ISR-002** | Israel | - | CP078843 |
| **HpGP-ISR-003** | Israel | + | CP078842 |
| **HpGP-ISR-004** | Israel | - | CP078841 |
| **HpGP-ISR-005** | Israel | - | CP078840 |
| **HpGP-ISR-006** | Israel | + | CP078839 |
| **HpGP-ISR-007** | Israel | - | CP078838 |
| **HpGP-ISR-008** | Israel | - | CP078837 |
| **HpGP-ISR-009** | Israel | - | CP078836 |
| **HpGP-ISR-010** | Israel | - | CP078835 |
| **HpGP-ITA-001** | Italy | - | CP079507 |
| **HpGP-ITA-002** | Italy | - | CP079506 |
| **HpGP-ITA-003** | Italy | + | CP079505 |
| **HpGP-ITA-004** | Italy | - | CP079504 |
| **HpGP-ITA-005** | Italy | - | CP079503 |
| **HpGP-ITA-006** | Italy | - | CP079502 |
| **HpGP-ITA-007** | Italy | + | CP079501 |
| **HpGP-ITA-008** | Italy | + | CP079500 |
| **HpGP-ITA-009** | Italy | + | CP079499 |
| **HpGP-ITA-010** | Italy | - | CP079498 |
| **HpGP-ITA-011** | Italy | - | CP079497 |
| **HpGP-ITA-012** | Italy | + | CP079496 |
| **HpGP-ITA-013** | Italy | - | CP079495 |
| **HpGP-ITA-014** | Italy | + | CP079494 |
| **HpGP-ITA-015** | Italy | + | CP079493 |
| **HpGP-ITA-016** | Italy | + | CP079492 |
| **HpGP-ITA-017** | Italy | - | CP079491 |
| **HpGP-ITA-018** | Italy | + | CP079490 |
| **HpGP-ITA-019** | Italy | + | CP079489 |
| **HpGP-ITA-020** | Italy | + | CP079488 |
| **HpGP-ITA-022** | Italy | + | CP079487 |
| **HpGP-ITA-023** | Italy | + | CP079486 |
| **HpGP-ITA-024** | Italy | + | CP079485 |
| **HpGP-ITA-025** | Italy | - | CP079484 |
| **HpGP-ITA-026** | Italy | + | CP079483 |
| **HpGP-ITA-027** | Italy | + | CP079482 |
| **HpGP-ITA-028** | Italy | + | CP079481 |
| **HpGP-ITA-029** | Italy | + | CP079480 |
| **HpGP-ITA-030** | Italy | + | CP079479 |
| **HpGP-JAP-002** | Japan | - | CP079478 |
| **HpGP-JAP-009** | Japan | + | CP079477 |
| **HpGP-JAP-010** | Japan | + | CP079476 |
| **HpGP-JAP-012** | Japan | + | CP079475 |
| **HpGP-JAP-013** | Japan | + | CP079474 |
| **HpGP-JAP-015** | Japan | + | CP079473 |
| **HpGP-JAP-016** | Japan | + | CP079472 |
| **HpGP-JAP-017** | Japan | + | CP079471 |
| **HpGP-JAP-019** | Japan | + | CP079470 |
| **HpGP-JAP-021** | Japan | + | CP079469 |
| **HpGP-JAP-022** | Japan | + | CP079468 |
| **HpGP-JAP-026** | Japan | + | CP079467 |
| **HpGP-JAP-028** | Japan | + | CP079466 |
| **HpGP-JAP-029** | Japan | + | CP079465 |
| **HpGP-JAP-101** | Japan | + | CP078834 |
| **HpGP-JAP-102** | Japan | + | CP078833 |
| **HpGP-JAP-103** | Japan | - | CP078832 |
| **HpGP-JAP-104** | Japan | + | CP078831 |
| **HpGP-JAP-105** | Japan | - | CP078830 |
| **HpGP-JAP-106** | Japan | - | CP078829 |
| **HpGP-JAP-107** | Japan | + | CP078828 |
| **HpGP-JAP-108** | Japan | + | CP078827 |
| **HpGP-JAP-109** | Japan | + | CP078826 |
| **HpGP-JAP-110** | Japan | + | CP078825 |
| **HpGP-JAP-111** | Japan | + | CP078824 |
| **HpGP-JAP-112** | Japan | + | CP078823 |
| **HpGP-JAP-113** | Japan | + | CP078822 |
| **HpGP-JAP-114** | Japan | + | CP078821 |
| **HpGP-JAP-115** | Japan | + | CP078820 |
| **HpGP-JOR-001** | Jordan | + | CP078819 |
| **HpGP-JOR-002** | Jordan | - | CP078818 |
| **HpGP-JOR-003** | Jordan | + | CP078817 |
| **HpGP-JOR-004** | Jordan | + | CP078816 |
| **HpGP-JOR-005** | Jordan | + | CP078815 |
| **HpGP-JOR-006** | Jordan | + | CP078814 |
| **HpGP-JOR-007** | Jordan | + | CP078813 |
| **HpGP-JOR-008** | Jordan | + | CP078812 |
| **HpGP-JOR-009** | Jordan | - | CP078811 |
| **HpGP-JOR-010** | Jordan | + | CP078810 |
| **HpGP-KAZ-001** | Kazakhstan | - | CP078809 |
| **HpGP-KAZ-002** | Kazakhstan | + | CP078808 |
| **HpGP-KGZ-001** | Kyrgyzstan | + | CP078807 |
| **HpGP-KGZ-002** | Kyrgyzstan | + | CP078806 |
| **HpGP-KGZ-003** | Kyrgyzstan | - | CP078805 |
| **HpGP-KGZ-004** | Kyrgyzstan | + | CP078804 |
| **HpGP-KGZ-005** | Kyrgyzstan | + | CP078803 |
| **HpGP-KGZ-006** | Kyrgyzstan | - | CP078802 |
| **HpGP-KGZ-007** | Kyrgyzstan | + | CP078801 |
| **HpGP-KGZ-008** | Kyrgyzstan | + | CP078800 |
| **HpGP-KGZ-009** | Kyrgyzstan | + | CP078799 |
| **HpGP-KGZ-010** | Kyrgyzstan | + | CP078798 |
| **HpGP-KOR-001** | South Korea | + | CP079464 |
| **HpGP-KOR-002** | South Korea | + | CP079463 |
| **HpGP-KOR-004** | South Korea | + | CP079462 |
| **HpGP-KOR-005** | South Korea | + | CP079461 |
| **HpGP-KOR-006** | South Korea | + | CP079460 |
| **HpGP-KOR-007** | South Korea | - | CP079459 |
| **HpGP-KOR-008** | South Korea | + | CP079458 |
| **HpGP-KOR-009** | South Korea | + | CP079457 |
| **HpGP-KOR-010** | South Korea | + | CP079456 |
| **HpGP-KOR-012** | South Korea | + | CP079455 |
| **HpGP-KOR-013** | South Korea | + | CP078797 |
| **HpGP-KOR-014** | South Korea | + | CP078796 |
| **HpGP-KOR-016** | South Korea | + | CP078795 |
| **HpGP-KOR-019** | South Korea | + | CP078794 |
| **HpGP-KOR-020** | South Korea | + | CP078793 |
| **HpGP-KOR-021** | South Korea | + | CP078792 |
| **HpGP-KOR-022** | South Korea | + | CP078791 |
| **HpGP-KOR-023** | South Korea | + | CP078790 |
| **HpGP-KOR-024** | South Korea | + | CP078789 |
| **HpGP-KOR-025** | South Korea | + | CP078788 |
| **HpGP-KOR-027** | South Korea | + | CP078787 |
| **HpGP-KOR-028** | South Korea | + | CP078786 |
| **HpGP-KOR-029** | South Korea | + | CP078785 |
| **HpGP-KOR-032** | South Korea | + | CP079454 |
| **HpGP-KOR-033** | South Korea | + | CP079453 |
| **HpGP-KOR-034** | South Korea | + | CP079452 |
| **HpGP-KOR-035** | South Korea | + | CP079451 |
| **HpGP-KOR-036** | South Korea | + | CP079450 |
| **HpGP-KOR-037** | South Korea | + | CP079449 |
| **HpGP-KOR-041** | South Korea | + | CP079448 |
| **HpGP-KOR-042** | South Korea | + | CP079447 |
| **HpGP-KOR-043** | South Korea | + | CP079446 |
| **HpGP-KOR-044** | South Korea | + | CP079445 |
| **HpGP-KOR-045** | South Korea | + | CP079444 |
| **HpGP-KOR-046** | South Korea | + | CP079443 |
| **HpGP-KOR-047** | South Korea | + | CP079442 |
| **HpGP-KOR-048** | South Korea | + | CP079441 |
| **HpGP-KOR-049** | South Korea | + | CP079440 |
| **HpGP-KOR-050** | South Korea | + | CP079439 |
| **HpGP-KOR-051** | South Korea | + | CP078784 |
| **HpGP-KOR-052** | South Korea | + | CP078783 |
| **HpGP-KOR-053** | South Korea | + | CP078782 |
| **HpGP-KOR-101** | South Korea | + | CP079438 |
| **HpGP-KOR-105** | South Korea | + | CP079437 |
| **HpGP-KOR-109** | South Korea | + | CP079436 |
| **HpGP-KOR-110** | South Korea | + | CP079435 |
| **HpGP-KOR-111** | South Korea | + | CP078781 |
| **HpGP-KOR-112** | South Korea | + | CP078780 |
| **HpGP-KOR-114** | South Korea | + | CP078778 |
| **HpGP-KOR-115** | South Korea | + | CP078777 |
| **HpGP-KOR-116** | South Korea | + | CP078776 |
| **HpGP-KOR-117** | South Korea | + | CP078504 |
| **HpGP-KOR-118** | South Korea | + | CP078503 |
| **HpGP-KOR-119** | South Korea | + | CP078502 |
| **HpGP-LAT-001** | Latvia | - | CP094499 |
| **HpGP-LAT-002** | Latvia | - | CP079433 |
| **HpGP-LAT-003** | Latvia | + | CP094498 |
| **HpGP-LAT-004** | Latvia | + | CP079431 |
| **HpGP-LAT-005** | Latvia | + | CP079430 |
| **HpGP-LAT-006** | Latvia | + | CP094497 |
| **HpGP-LAT-007** | Latvia | + | CP079428 |
| **HpGP-LAT-008** | Latvia | - | CP079427 |
| **HpGP-LAT-009** | Latvia | + | CP079426 |
| **HpGP-LAT-010** | Latvia | - | CP079425 |
| **HpGP-LAT-011** | Latvia | + | CP078501 |
| **HpGP-LAT-012** | Latvia | + | CP079424 |
| **HpGP-LAT-014** | Latvia | + | CP079422 |
| **HpGP-LAT-015** | Latvia | + | CP079421 |
| **HpGP-LAT-017** | Latvia | + | CP079420 |
| **HpGP-LAT-018** | Latvia | + | CP094496 |
| **HpGP-LAT-019** | Latvia | + | CP079418 |
| **HpGP-LAT-020** | Latvia | + | CP079417 |
| **HpGP-LAT-021** | Latvia | + | CP079416 |
| **HpGP-LAT-022** | Latvia | + | CP079415 |
| **HpGP-LAT-023** | Latvia | + | CP079414 |
| **HpGP-LAT-024** | Latvia | + | CP079413 |
| **HpGP-LAT-025** | Latvia | + | CP079412 |
| **HpGP-LAT-026** | Latvia | + | CP079411 |
| **HpGP-LAT-027** | Latvia | + | CP079410 |
| **HpGP-LAT-028** | Latvia | + | CP079409 |
| **HpGP-LAT-029** | Latvia | + | CP079408 |
| **HpGP-LAT-030** | Latvia | + | CP079407 |
| **HpGP-LAT-031** | Latvia | + | CP078500 |
| **HpGP-LAT-032** | Latvia | + | CP078499 |
| **HpGP-LAT-033** | Latvia | - | CP078498 |
| **HpGP-LAT-034** | Latvia | + | CP078497 |
| **HpGP-LAT-035** | Latvia | + | CP078496 |
| **HpGP-LAT-036** | Latvia | + | CP078495 |
| **HpGP-LIT-001** | Lithuania | - | CP079406 |
| **HpGP-LIT-002** | Lithuania | - | CP079405 |
| **HpGP-LIT-003** | Lithuania | + | CP079404 |
| **HpGP-LIT-004** | Lithuania | + | CP079403 |
| **HpGP-LIT-006** | Lithuania | + | CP079402 |
| **HpGP-LIT-007** | Lithuania | + | CP079401 |
| **HpGP-LIT-008** | Lithuania | - | CP079400 |
| **HpGP-LIT-010** | Lithuania | + | CP079399 |
| **HpGP-LIT-012** | Lithuania | - | CP079398 |
| **HpGP-LIT-013** | Lithuania | + | CP079397 |
| **HpGP-LIT-020** | Lithuania | + | CP079396 |
| **HpGP-LIT-021** | Lithuania | + | CP078494 |
| **HpGP-LIT-023** | Lithuania | + | CP079395 |
| **HpGP-LIT-024** | Lithuania | + | CP079394 |
| **HpGP-LIT-025** | Lithuania | - | CP079393 |
| **HpGP-LIT-026** | Lithuania | + | CP078493 |
| **HpGP-LIT-028** | Lithuania | + | CP078492 |
| **HpGP-LIT-029** | Lithuania | + | CP079392 |
| **HpGP-LIT-030** | Lithuania | + | CP079391 |
| **HpGP-LIT-031** | Lithuania | + | CP078491 |
| **HpGP-LIT-032** | Lithuania | + | CP078490 |
| **HpGP-LIT-035** | Lithuania | + | CP079390 |
| **HpGP-LIT-041** | Lithuania | - | CP078489 |
| **HpGP-MAL-002** | Malaysia | + | CP079388 |
| **HpGP-MAL-003** | Malaysia | + | CP079387 |
| **HpGP-MAL-004** | Malaysia | + | CP079386 |
| **HpGP-MAL-005** | Malaysia | + | CP079385 |
| **HpGP-MAL-006** | Malaysia | + | CP079384 |
| **HpGP-MAL-007** | Malaysia | + | CP079383 |
| **HpGP-MAL-008** | Malaysia | + | CP079382 |
| **HpGP-MAL-010** | Malaysia | + | CP079381 |
| **HpGP-MAL-012** | Malaysia | + | CP079380 |
| **HpGP-MAL-015** | Malaysia | + | CP079379 |
| **HpGP-MAL-016** | Malaysia | + | CP079378 |
| **HpGP-MAL-017** | Malaysia | + | CP079377 |
| **HpGP-MAL-018** | Malaysia | + | CP079376 |
| **HpGP-MAL-019** | Malaysia | + | CP079375 |
| **HpGP-MAL-020** | Malaysia | + | CP079374 |
| **HpGP-MAL-021** | Malaysia | - | CP079373 |
| **HpGP-MAL-022** | Malaysia | + | CP079372 |
| **HpGP-MAL-023** | Malaysia | + | CP079371 |
| **HpGP-MAL-025** | Malaysia | + | CP079370 |
| **HpGP-MEX-001** | Mexico | + | CP079369 |
| **HpGP-MEX-002** | Mexico | + | CP079368 |
| **HpGP-MEX-003** | Mexico | + | CP079367 |
| **HpGP-MEX-004** | Mexico | + | CP079366 |
| **HpGP-MEX-006** | Mexico | + | CP079365 |
| **HpGP-MEX-007** | Mexico | - | CP079364 |
| **HpGP-MEX-008** | Mexico | - | CP079363 |
| **HpGP-MEX-009** | Mexico | - | CP079362 |
| **HpGP-MEX-010** | Mexico | + | CP079361 |
| **HpGP-MEX-012** | Mexico | + | CP079360 |
| **HpGP-MEX-023** | Mexico | - | CP079359 |
| **HpGP-MEX-024** | Mexico | + | CP079358 |
| **HpGP-MEX-027** | Mexico | + | CP079357 |
| **HpGP-MEX-028** | Mexico | - | CP079356 |
| **HpGP-MEX-032** | Mexico | + | CP079355 |
| **HpGP-MEX-035** | Mexico | + | CP079354 |
| **HpGP-MEX-040** | Mexico | + | CP079353 |
| **HpGP-MEX-041** | Mexico | + | CP079352 |
| **HpGP-MEX-043** | Mexico | + | CP079351 |
| **HpGP-MEX-044** | Mexico | + | CP079350 |
| **HpGP-MEX-045** | Mexico | + | CP078488 |
| **HpGP-MEX-046** | Mexico | + | CP078487 |
| **HpGP-MMR-001** | Myanmar | + | CP078486 |
| **HpGP-MMR-002** | Myanmar | + | CP078485 |
| **HpGP-MMR-003** | Myanmar | + | CP078484 |
| **HpGP-MMR-004** | Myanmar | + | CP078483 |
| **HpGP-MMR-005** | Myanmar | + | CP078482 |
| **HpGP-MMR-006** | Myanmar | + | CP078481 |
| **HpGP-MMR-007** | Myanmar | + | CP078480 |
| **HpGP-MMR-008** | Myanmar | + | CP078479 |
| **HpGP-MMR-009** | Myanmar | + | CP078478 |
| **HpGP-MMR-010** | Myanmar | + | CP078477 |
| **HpGP-MMR-011** | Myanmar | + | CP078476 |
| **HpGP-MMR-012** | Myanmar | + | CP078475 |
| **HpGP-NGA-006** | Nigeria | + | CP078474 |
| **HpGP-NGA-010** | Nigeria | + | CP078473 |
| **HpGP-NGA-017** | Nigeria | + | CP078472 |
| **HpGP-NGA-019** | Nigeria | + | CP078471 |
| **HpGP-NPL-001** | Nepal | - | CP078470 |
| **HpGP-NPL-002** | Nepal | - | CP078469 |
| **HpGP-NPL-003** | Nepal | + | CP078468 |
| **HpGP-NPL-004** | Nepal | + | CP078467 |
| **HpGP-NPL-005** | Nepal | + | CP078466 |
| **HpGP-NPL-006** | Nepal | + | CP078465 |
| **HpGP-NPL-007** | Nepal | + | CP078464 |
| **HpGP-NPL-008** | Nepal | + | CP078463 |
| **HpGP-NPL-009** | Nepal | + | CP078462 |
| **HpGP-NPL-010** | Nepal | + | CP078461 |
| **HpGP-NPL-011** | Nepal | + | CP078460 |
| **HpGP-NPL-012** | Nepal | + | CP078459 |
| **HpGP-NPL-013** | Nepal | + | CP078458 |
| **HpGP-PER-011** | Peru | + | CP079349 |
| **HpGP-PER-013** | Peru | + | CP079348 |
| **HpGP-PER-014** | Peru | + | CP079347 |
| **HpGP-PER-016** | Peru | + | CP079345 |
| **HpGP-PER-017** | Peru | + | CP079344 |
| **HpGP-PER-018** | Peru | + | CP079343 |
| **HpGP-PER-019** | Peru | + | CP079342 |
| **HpGP-PER-020** | Peru | + | CP079341 |
| **HpGP-PER-021** | Peru | + | CP079340 |
| **HpGP-PER-101** | Peru | + | CP078457 |
| **HpGP-PER-102** | Peru | + | CP078456 |
| **HpGP-PER-103** | Peru | - | CP078455 |
| **HpGP-PER-104** | Peru | + | CP078454 |
| **HpGP-PER-105** | Peru | + | CP078453 |
| **HpGP-PER-106** | Peru | + | CP078452 |
| **HpGP-PER-107** | Peru | + | CP078451 |
| **HpGP-PER-108** | Peru | + | CP078450 |
| **HpGP-PER-109** | Peru | + | CP078449 |
| **HpGP-PER-110** | Peru | + | CP078448 |
| **HpGP-PER-111** | Peru | + | CP078447 |
| **HpGP-PER-112** | Peru | + | CP078446 |
| **HpGP-PER-113** | Peru | + | CP078445 |
| **HpGP-PER-114** | Peru | + | CP078444 |
| **HpGP-PER-115** | Peru | + | CP078443 |
| **HpGP-PER-116** | Peru | + | CP078442 |
| **HpGP-PER-117** | Peru | + | CP078441 |
| **HpGP-PER-119** | Peru | + | CP078440 |
| **HpGP-PER-121** | Peru | + | CP078439 |
| **HpGP-PER-124** | Peru | + | CP078438 |
| **HpGP-PER-130** | Peru | + | CP078437 |
| **HpGP-PER-136** | Peru | + | CP078436 |
| **HpGP-PER-138** | Peru | + | CP078435 |
| **HpGP-PER-140** | Peru | + | CP078434 |
| **HpGP-POL-001** | Poland | + | CP078433 |
| **HpGP-POL-002** | Poland | - | CP078432 |
| **HpGP-POL-003** | Poland | - | CP078431 |
| **HpGP-POL-004** | Poland | - | CP078430 |
| **HpGP-POL-005** | Poland | + | CP078429 |
| **HpGP-POL-006** | Poland | - | CP078428 |
| **HpGP-POL-007** | Poland | + | CP078427 |
| **HpGP-POL-008** | Poland | - | CP078426 |
| **HpGP-POL-009** | Poland | + | CP078425 |
| **HpGP-POL-010** | Poland | - | CP078424 |
| **HpGP-POL-101** | Poland | - | CP078423 |
| **HpGP-POL-102** | Poland | + | CP078422 |
| **HpGP-POL-103** | Poland | + | CP078421 |
| **HpGP-POL-104** | Poland | + | CP078420 |
| **HpGP-POL-105** | Poland | - | CP078419 |
| **HpGP-POL-106** | Poland | - | CP078418 |
| **HpGP-POL-107** | Poland | - | CP078417 |
| **HpGP-POL-108** | Poland | - | CP078416 |
| **HpGP-POL-109** | Poland | - | CP078415 |
| **HpGP-POL-110** | Poland | - | CP078414 |
| **HpGP-POR-001** | Portugal | - | CP079339 |
| **HpGP-POR-002** | Portugal | + | CP079338 |
| **HpGP-POR-003** | Portugal | - | CP079337 |
| **HpGP-POR-004** | Portugal | + | CP079336 |
| **HpGP-POR-005** | Portugal | + | CP079335 |
| **HpGP-POR-006** | Portugal | + | CP079334 |
| **HpGP-POR-007** | Portugal | - | CP079333 |
| **HpGP-POR-008** | Portugal | - | CP079332 |
| **HpGP-POR-009** | Portugal | - | CP079331 |
| **HpGP-POR-010** | Portugal | - | CP079290 |
| **HpGP-POR-011** | Portugal | - | CP079330 |
| **HpGP-POR-012** | Portugal | + | CP079329 |
| **HpGP-POR-013** | Portugal | - | CP079328 |
| **HpGP-POR-015** | Portugal | - | CP079327 |
| **HpGP-POR-018** | Portugal | - | CP079325 |
| **HpGP-POR-101** | Portugal | - | CP078413 |
| **HpGP-POR-102** | Portugal | - | CP078412 |
| **HpGP-POR-103** | Portugal | - | CP078411 |
| **HpGP-POR-104** | Portugal | + | CP078410 |
| **HpGP-POR-105** | Portugal | - | CP078409 |
| **HpGP-POR-106** | Portugal | - | CP078408 |
| **HpGP-POR-107** | Portugal | - | CP078407 |
| **HpGP-POR-108** | Portugal | - | CP078406 |
| **HpGP-POR-109** | Portugal | - | CP078405 |
| **HpGP-POR-110** | Portugal | + | CP078404 |
| **HpGP-POR-111** | Portugal | - | CP078403 |
| **HpGP-POR-112** | Portugal | + | CP078402 |
| **HpGP-POR-113** | Portugal | + | CP078401 |
| **HpGP-POR-114** | Portugal | + | CP078400 |
| **HpGP-POR-115** | Portugal | + | CP078399 |
| **HpGP-PRI-001** | Puerto Rico | - | CP078398 |
| **HpGP-PRI-002** | Puerto Rico | - | CP078397 |
| **HpGP-PRI-004** | Puerto Rico | - | CP078396 |
| **HpGP-PRI-005** | Puerto Rico | - | CP078395 |
| **HpGP-PRI-006** | Puerto Rico | - | CP078394 |
| **HpGP-PRI-007** | Puerto Rico | - | CP078393 |
| **HpGP-PRI-008** | Puerto Rico | + | CP078392 |
| **HpGP-RUS-001** | Russian Federation | + | CP078390 |
| **HpGP-RUS-002** | Russian Federation | - | CP078389 |
| **HpGP-RUS-003** | Russian Federation | + | CP078388 |
| **HpGP-RUS-004** | Russian Federation | + | CP078387 |
| **HpGP-RUS-006** | Russian Federation | - | CP078386 |
| **HpGP-RUS-007** | Russian Federation | + | CP078385 |
| **HpGP-RUS-008** | Russian Federation | - | CP078384 |
| **HpGP-RUS-009** | Russian Federation | + | CP078383 |
| **HpGP-RUS-010** | Russian Federation | + | CP078382 |
| **HpGP-RUS-011** | Russian Federation | - | CP078381 |
| **HpGP-SGP-002** | Singapore | - | CP078380 |
| **HpGP-SGP-003** | Singapore | + | CP078379 |
| **HpGP-SGP-004** | Singapore | + | CP078378 |
| **HpGP-SGP-005** | Singapore | + | CP078377 |
| **HpGP-SGP-006** | Singapore | + | CP078376 |
| **HpGP-SGP-007** | Singapore | + | CP078375 |
| **HpGP-SGP-008** | Singapore | + | CP078374 |
| **HpGP-SGP-009** | Singapore | + | CP078373 |
| **HpGP-SGP-012** | Singapore | + | CP078372 |
| **HpGP-SGP-013** | Singapore | + | CP078371 |
| **HpGP-SGP-014** | Singapore | + | CP078370 |
| **HpGP-SGP-015** | Singapore | + | CP078369 |
| **HpGP-SGP-017** | Singapore | + | CP078368 |
| **HpGP-SGP-018** | Singapore | + | CP078367 |
| **HpGP-SGP-019** | Singapore | + | CP078366 |
| **HpGP-SGP-024** | Singapore | + | CP078365 |
| **HpGP-SGP-026** | Singapore | + | CP078364 |
| **HpGP-SGP-032** | Singapore | + | CP078363 |
| **HpGP-SGP-033** | Singapore | + | CP078362 |
| **HpGP-SGP-036** | Singapore | + | CP078361 |
| **HpGP-SGP-038** | Singapore | + | CP078360 |
| **HpGP-SPA-001** | Spain | - | CP079324 |
| **HpGP-SPA-002** | Spain | - | CP079323 |
| **HpGP-SPA-003** | Spain | - | CP079322 |
| **HpGP-SPA-004** | Spain | + | CP079321 |
| **HpGP-SPA-005** | Spain | - | CP079320 |
| **HpGP-SPA-006** | Spain | - | CP079319 |
| **HpGP-SPA-007** | Spain | - | CP079318 |
| **HpGP-SPA-008** | Spain | - | CP079317 |
| **HpGP-SPA-009** | Spain | - | CP079316 |
| **HpGP-SPA-010** | Spain | - | CP079315 |
| **HpGP-SPA-011** | Spain | + | CP079314 |
| **HpGP-SPA-012** | Spain | - | CP079313 |
| **HpGP-SPA-013** | Spain | + | CP079312 |
| **HpGP-SPA-014** | Spain | + | CP079311 |
| **HpGP-SPA-015** | Spain | + | CP079310 |
| **HpGP-SPA-016** | Spain | + | CP079309 |
| **HpGP-SPA-017** | Spain | + | CP079308 |
| **HpGP-SPA-018** | Spain | + | CP079307 |
| **HpGP-SPA-019** | Spain | + | CP079306 |
| **HpGP-SPA-020** | Spain | + | CP079305 |
| **HpGP-SPA-101** | Spain | - | CP078359 |
| **HpGP-SPA-102** | Spain | - | CP078358 |
| **HpGP-SPA-103** | Spain | - | CP078357 |
| **HpGP-SPA-104** | Spain | - | CP078356 |
| **HpGP-SPA-105** | Spain | - | CP078355 |
| **HpGP-SPA-107** | Spain | - | CP078354 |
| **HpGP-SPA-108** | Spain | + | CP078353 |
| **HpGP-SPA-110** | Spain | - | CP078352 |
| **HpGP-SPA-111** | Spain | - | CP078351 |
| **HpGP-SPA-201** | Spain | + | CP078350 |
| **HpGP-SPA-202** | Spain | + | CP078349 |
| **HpGP-SPA-203** | Spain | - | CP078348 |
| **HpGP-SPA-204** | Spain | - | CP078347 |
| **HpGP-SPA-205** | Spain | - | CP078346 |
| **HpGP-SPA-206** | Spain | - | CP078345 |
| **HpGP-SPA-207** | Spain | + | CP078344 |
| **HpGP-SPA-209** | Spain | - | CP078343 |
| **HpGP-SPA-210** | Spain | - | CP078342 |
| **HpGP-SPA-301** | Spain | + | CP078341 |
| **HpGP-SPA-302** | Spain | - | CP078340 |
| **HpGP-SPA-303** | Spain | - | CP078339 |
| **HpGP-SPA-304** | Spain | - | CP078338 |
| **HpGP-SPA-305** | Spain | + | CP078337 |
| **HpGP-SPA-306** | Spain | - | CP078336 |
| **HpGP-SPA-307** | Spain | - | CP078335 |
| **HpGP-SPA-308** | Spain | - | CP078334 |
| **HpGP-SPA-309** | Spain | + | CP078333 |
| **HpGP-SPA-310** | Spain | - | CP078332 |
| **HpGP-SPA-311** | Spain | + | CP078331 |
| **HpGP-SPA-312** | Spain | + | CP078330 |
| **HpGP-SPA-313** | Spain | + | CP078329 |
| **HpGP-SPA-314** | Spain | + | CP078328 |
| **HpGP-SPA-315** | Spain | + | CP078327 |
| **HpGP-SPA-316** | Spain | - | CP078326 |
| **HpGP-SPA-317** | Spain | + | CP078325 |
| **HpGP-SPA-318** | Spain | + | CP078324 |
| **HpGP-SPA-319** | Spain | + | CP078323 |
| **HpGP-SPA-320** | Spain | + | CP078322 |
| **HpGP-SPA-321** | Spain | + | CP078321 |
| **HpGP-SPA-322** | Spain | + | CP078320 |
| **HpGP-SPA-323** | Spain | + | CP078319 |
| **HpGP-SPA-324** | Spain | - | CP078318 |
| **HpGP-SPA-325** | Spain | + | CP078317 |
| **HpGP-SPA-401** | Spain | - | CP078316 |
| **HpGP-SPA-402** | Spain | + | CP078315 |
| **HpGP-SPA-403** | Spain | + | CP078314 |
| **HpGP-SPA-404** | Spain | + | CP078313 |
| **HpGP-SPA-405** | Spain | + | CP078312 |
| **HpGP-SPA-406** | Spain | - | CP078311 |
| **HpGP-SPA-407** | Spain | - | CP078310 |
| **HpGP-SPA-409** | Spain | + | CP078309 |
| **HpGP-SPA-413** | Spain | - | CP078308 |
| **HpGP-SPA-414** | Spain | - | CP078307 |
| **HpGP-SPA-501** | Spain | - | CP078306 |
| **HpGP-SPA-502** | Spain | - | CP078305 |
| **HpGP-SPA-503** | Spain | - | CP078304 |
| **HpGP-SPA-504** | Spain | + | CP078303 |
| **HpGP-SPA-505** | Spain | - | CP078302 |
| **HpGP-SPA-506** | Spain | - | CP078301 |
| **HpGP-SPA-507** | Spain | + | CP078300 |
| **HpGP-SPA-508** | Spain | - | CP078299 |
| **HpGP-SPA-509** | Spain | - | CP078298 |
| **HpGP-SPA-510** | Spain | - | CP078297 |
| **HpGP-SPA-601** | Spain | + | CP078296 |
| **HpGP-SPA-604** | Spain | - | CP078295 |
| **HpGP-SPA-606** | Spain | + | CP078294 |
| **HpGP-SPA-607** | Spain | - | CP078293 |
| **HpGP-SPA-609** | Spain | - | CP078292 |
| **HpGP-SPA-610** | Spain | - | CP078291 |
| **HpGP-SPA-611** | Spain | + | CP078290 |
| **HpGP-SPA-701** | Spain | - | CP078289 |
| **HpGP-SPA-702** | Spain | - | CP078288 |
| **HpGP-SPA-703** | Spain | - | CP078287 |
| **HpGP-SPA-704** | Spain | - | CP078286 |
| **HpGP-SPA-705** | Spain | - | CP078285 |
| **HpGP-SPA-706** | Spain | + | CP078284 |
| **HpGP-SPA-707** | Spain | - | CP078283 |
| **HpGP-SPA-709** | Spain | + | CP078282 |
| **HpGP-SPA-710** | Spain | - | CP078281 |
| **HpGP-SPA-801** | Spain | + | CP078280 |
| **HpGP-SPA-802** | Spain | - | CP078279 |
| **HpGP-SPA-803** | Spain | - | CP078278 |
| **HpGP-SPA-804** | Spain | - | CP078277 |
| **HpGP-SPA-805** | Spain | - | CP078276 |
| **HpGP-SPA-806** | Spain | - | CP078275 |
| **HpGP-SPA-807** | Spain | + | CP078274 |
| **HpGP-SWE-001** | Sweden | - | CP079304 |
| **HpGP-SWE-002** | Sweden | + | CP079303 |
| **HpGP-SWE-003** | Sweden | + | CP079302 |
| **HpGP-SWE-004** | Sweden | + | CP079301 |
| **HpGP-SWE-005** | Sweden | - | CP079300 |
| **HpGP-SWE-006** | Sweden | - | CP079299 |
| **HpGP-SWE-007** | Sweden | + | CP079298 |
| **HpGP-SWE-008** | Sweden | + | CP079297 |
| **HpGP-SWE-010** | Sweden | + | CP079296 |
| **HpGP-SWE-012** | Sweden | + | CP079295 |
| **HpGP-SWE-013** | Sweden | + | CP078273 |
| **HpGP-SWE-014** | Sweden | + | CP079294 |
| **HpGP-SWE-015** | Sweden | - | CP079293 |
| **HpGP-SWE-016** | Sweden | + | CP079292 |
| **HpGP-SWE-017** | Sweden | + | CP079291 |
| **HpGP-SWE-018** | Sweden | + | CP079289 |
| **HpGP-SWE-019** | Sweden | + | CP079288 |
| **HpGP-SWE-021** | Sweden | - | CP079287 |
| **HpGP-SWE-022** | Sweden | + | CP079286 |
| **HpGP-SWE-024** | Sweden | + | CP079285 |
| **HpGP-SWE-025** | Sweden | + | CP079284 |
| **HpGP-SWE-026** | Sweden | + | CP079283 |
| **HpGP-SWE-027** | Sweden | + | CP079282 |
| **HpGP-SWE-028** | Sweden | + | CP079281 |
| **HpGP-SWE-030** | Sweden | + | CP079280 |
| **HpGP-SWE-031** | Sweden | + | CP079279 |
| **HpGP-SWE-032** | Sweden | + | CP079278 |
| **HpGP-SWE-033** | Sweden | - | CP079277 |
| **HpGP-SWE-034** | Sweden | + | CP079276 |
| **HpGP-SWE-036** | Sweden | + | CP079275 |
| **HpGP-SWT-001** | Switzerland | - | CP079274 |
| **HpGP-SWT-002** | Switzerland | - | CP079273 |
| **HpGP-SWT-003** | Switzerland | - | CP087592 |
| **HpGP-SWT-004** | Switzerland | + | CP079271 |
| **HpGP-SWT-005** | Switzerland | + | CP079270 |
| **HpGP-SWT-006** | Switzerland | - | CP079269 |
| **HpGP-SWT-007** | Switzerland | + | CP079268 |
| **HpGP-SWT-008** | Switzerland | + | CP079267 |
| **HpGP-SWT-010** | Switzerland | + | CP079266 |
| **HpGP-SWT-011** | Switzerland | - | CP079265 |
| **HpGP-SWT-012** | Switzerland | - | CP079264 |
| **HpGP-SWT-013** | Switzerland | - | CP079263 |
| **HpGP-SWT-014** | Switzerland | + | CP079262 |
| **HpGP-SWT-015** | Switzerland | - | CP079261 |
| **HpGP-SWT-017** | Switzerland | + | CP079260 |
| **HpGP-TUR-001** | Türkiye | + | CP078272 |
| **HpGP-TUR-002** | Türkiye | - | CP078271 |
| **HpGP-TUR-003** | Türkiye | - | CP078270 |
| **HpGP-TUR-004** | Türkiye | + | CP078269 |
| **HpGP-TUR-005** | Türkiye | - | CP078268 |
| **HpGP-TUR-006** | Türkiye | + | CP078267 |
| **HpGP-TUR-007** | Türkiye | - | CP078266 |
| **HpGP-TUR-008** | Türkiye | + | CP078265 |
| **HpGP-TUR-009** | Türkiye | - | CP078264 |
| **HpGP-TUR-010** | Türkiye | - | CP078263 |
| **HpGP-TUR-011** | Türkiye | - | CP078262 |
| **HpGP-TUR-014** | Türkiye | + | CP078258 |
| **HpGP-TUR-016** | **Türkiye** | **+** | **CP078257** |
| **HpGP-TUR-017** | **Türkiye** | **+** | **CP078256** |
| **HpGP-TUR-021** | **Türkiye** | **-** | **CP078255** |
| **HpGP-TUR-022** | **Türkiye** | **-** | **CP078254** |
| **HpGP-TUR-023** | **Türkiye** | **+** | **CP078253** |
| **HpGP-TWN-001** | **Taiwan** | **+** | **CP079259** |
| **HpGP-TWN-002** | **Taiwan** | **-** | **CP079258** |
| **HpGP-TWN-003** | **Taiwan** | **+** | **CP079257** |
| **HpGP-TWN-004** | **Taiwan** | **+** | **CP079256** |
| **HpGP-TWN-005** | **Taiwan** | **+** | **CP079255** |
| **HpGP-TWN-006** | **Taiwan** | **+** | **CP079254** |
| **HpGP-TWN-007** | **Taiwan** | **+** | **CP079253** |
| **HpGP-TWN-008** | **Taiwan** | **+** | **CP079252** |
| **HpGP-TWN-009** | **Taiwan** | **+** | **CP079251** |
| **HpGP-TWN-010** | **Taiwan** | **+** | **CP079250** |
| **HpGP-TWN-016** | **Taiwan** | **+** | **CP079249** |
| **HpGP-TWN-017** | **Taiwan** | **+** | **CP079248** |
| **HpGP-TWN-018** | **Taiwan** | **+** | **CP079247** |
| **HpGP-TWN-019** | **Taiwan** | **+** | **CP079246** |
| **HpGP-TWN-020** | **Taiwan** | **+** | **CP079245** |
| **HpGP-TWN-021** | **Taiwan** | **+** | **CP079244** |
| **HpGP-TWN-022** | **Taiwan** | **+** | **CP079243** |
| **HpGP-TWN-023** | **Taiwan** | **+** | **CP079242** |
| **HpGP-TWN-024** | **Taiwan** | **+** | **CP079241** |
| **HpGP-TWN-026** | **Taiwan** | **+** | **CP078252** |
| **HpGP-TWN-027** | **Taiwan** | **+** | **CP078251** |
| **HpGP-TWN-028** | **Taiwan** | **+** | **CP078250** |
| **HpGP-TWN-029** | **Taiwan** | **+** | **CP078249** |
| **HpGP-TWN-030** | **Taiwan** | **+** | **CP078248** |
| **HpGP-USA-001** | **USA** | **+** | **CP078247** |
| **HpGP-USA-002** | **USA** | **+** | **CP078246** |
| **HpGP-USA-003** | **USA** | **+** | **CP078245** |
| **HpGP-USA-004** | **USA** | **+** | **CP078244** |
| **HpGP-USA-005** | **USA** | **-** | **CP078243** |
| **HpGP-USA-008** | **USA** | **-** | **CP078242** |
| **HpGP-USA-009** | **USA** | **+** | **CP078241** |
| **HpGP-USA-011** | **USA** | **+** | **CP078240** |
| **HpGP-USA-101** | **USA** | **-** | **CP078239** |
| **HpGP-USA-102** | **USA** | **+** | **CP078238** |
| **HpGP-USA-104** | **USA** | **+** | **CP078237** |
| **HpGP-USA-105** | USA | + | CP078236 |
| **HpGP-USA-106** | USA | - | CP078235 |
| **HpGP-USA-115** | USA | - | CP078234 |
| **HpGP-USA-116** | USA | - | CP078233 |
| **HpGP-USA-119** | USA | - | CP078232 |
| **HpGP-USA-120** | USA | - | CP078231 |
| **HpGP-USA-123** | USA | - | CP078230 |
| **HpGP-USA-201** | USA | - | CP078229 |
| **HpGP-USA-202** | USA | + | CP078228 |
| **HpGP-USA-203** | USA | + | CP078227 |
| **HpGP-USA-204** | USA | + | CP078226 |
| **HpGP-USA-205** | USA | - | CP078225 |
| **HpGP-USA-206** | USA | + | CP078224 |
| **HpGP-USA-207** | USA | - | CP078223 |
| **HpGP-USA-208** | USA | - | CP078222 |
| **HpGP-USA-301** | USA | + | CP078221 |
| **HpGP-USA-302** | USA | + | CP078220 |
| **HpGP-USA-303** | USA | - | CP078219 |
| **HpGP-USA-304** | USA | + | CP078218 |
| **HpGP-USA-305** | USA | + | CP078217 |
| **HpGP-USA-306** | USA | + | CP078216 |
| **HpGP-USA-311** | USA | + | CP078215 |
| **HpGP-USA-401** | USA | - | CP078214 |
| **HpGP-USA-402** | USA | - | CP078213 |
| **HpGP-USA-403** | USA | + | CP078212 |
| **HpGP-USA-404** | USA | - | CP078211 |
| **HpGP-USA-405** | USA | - | CP078210 |
| **HpGP-USA-406** | USA | + | CP078209 |
| **HpGP-USA-407** | USA | - | CP078208 |
| **HpGP-USA-408** | USA | + | CP082186 |
| **HpGP-USA-409** | USA | + | CP078207 |
| **HpGP-USA-410** | USA | - | CP078206 |
| **HpGP-USA-411** | USA | - | CP078205 |
| **HpGP-USA-412** | USA | - | CP078204 |
| **HpGP-USA-413** | USA | + | CP078203 |
| **HpGP-USA-414** | USA | - | CP078202 |
| **HpGP-USA-415** | USA | + | CP078201 |
| **HpGP-USA-416** | USA | + | CP078200 |
| **HpGP-USA-417** | USA | + | CP078199 |
| **HpGP-USA-418** | USA | + | CP078198 |
| **HpGP-USA-419** | USA | + | CP078197 |
| **HpGP-USA-420** | USA | - | CP078196 |
| **HpGP-USA-421** | USA | + | CP078195 |
| **HpGP-USA-422** | USA | + | CP078194 |
| **HpGP-USA-423** | USA | - | CP078193 |
| **HpGP-USA-424** | USA | + | CP078192 |
| **HpGP-USA-425** | USA | + | CP078191 |
| **HpGP-USA-427** | USA | + | CP078190 |
| **HpGP-USA-428** | USA | + | CP078189 |
| **HpGP-USA-429** | USA | + | CP082185 |
| **HpGP-USA-431** | USA | - | CP078188 |
| **HpGP-USA-432** | USA | + | CP078187 |
| **HpGP-USA-433** | USA | + | CP078186 |
| **HpGP-USA-434** | USA | + | CP078185 |
| **HpGP-USA-435** | USA | - | CP078184 |
| **HpGP-USA-436** | USA | - | CP078183 |
| **HpGP-USA-437** | USA | + | CP078182 |
| **HpGP-VNM-001** | Vietnam | + | CP078181 |
| **HpGP-VNM-002** | Vietnam | + | CP078180 |
| **HpGP-VNM-003** | Vietnam | + | CP078179 |
| **HpGP-VNM-004** | Vietnam | + | CP078178 |
| **HpGP-VNM-005** | Vietnam | + | CP078177 |
| **HpGP-VNM-006** | Vietnam | + | CP078176 |
| **HpGP-VNM-007** | Vietnam | + | CP078175 |
| **HpGP-VNM-008** | Vietnam | + | CP078174 |
| **HpGP-VNM-009** | Vietnam | + | CP078173 |
| **HpGP-ZAF-001** | South Africa | - | CP078172 |
| **HpGP-ZAF-002** | South Africa | + | CP078171 |
| **HpGP-ZAF-004** | South Africa | + | CP078170 |
| **HpGP-ZAF-006** | South Africa | - | CP078169 |
| **HpGP-ZAF-007** | South Africa | - | CP078168 |
| **HpGP-ZAF-008** | South Africa | - | CP078167 |
| **HpGP-ZAF-009** | South Africa | - | CP078166 |
| **HpGP-ZAF-010** | South Africa | - | CP078165 |
| **J166_GC** | Europe | + | JQ685133 |
| **J99_derivate(IM)** | Europe | + | SRX8050361 |
| **mOut1** | Europe | + | JQ685134 |
| **mOut2** | Europe | + | JQ685135 |
| **mOut3** | Europe | + | JQ685136 |
| **mOut4** | Europe | + | JQ685137 |
| **rOut1** | Europe | + | JQ685138 |
| **rOut2** | Europe | + | JQ685139 |
| **rOut3** | Europe | + | JQ685140 |

**Supplementary Table 2.**
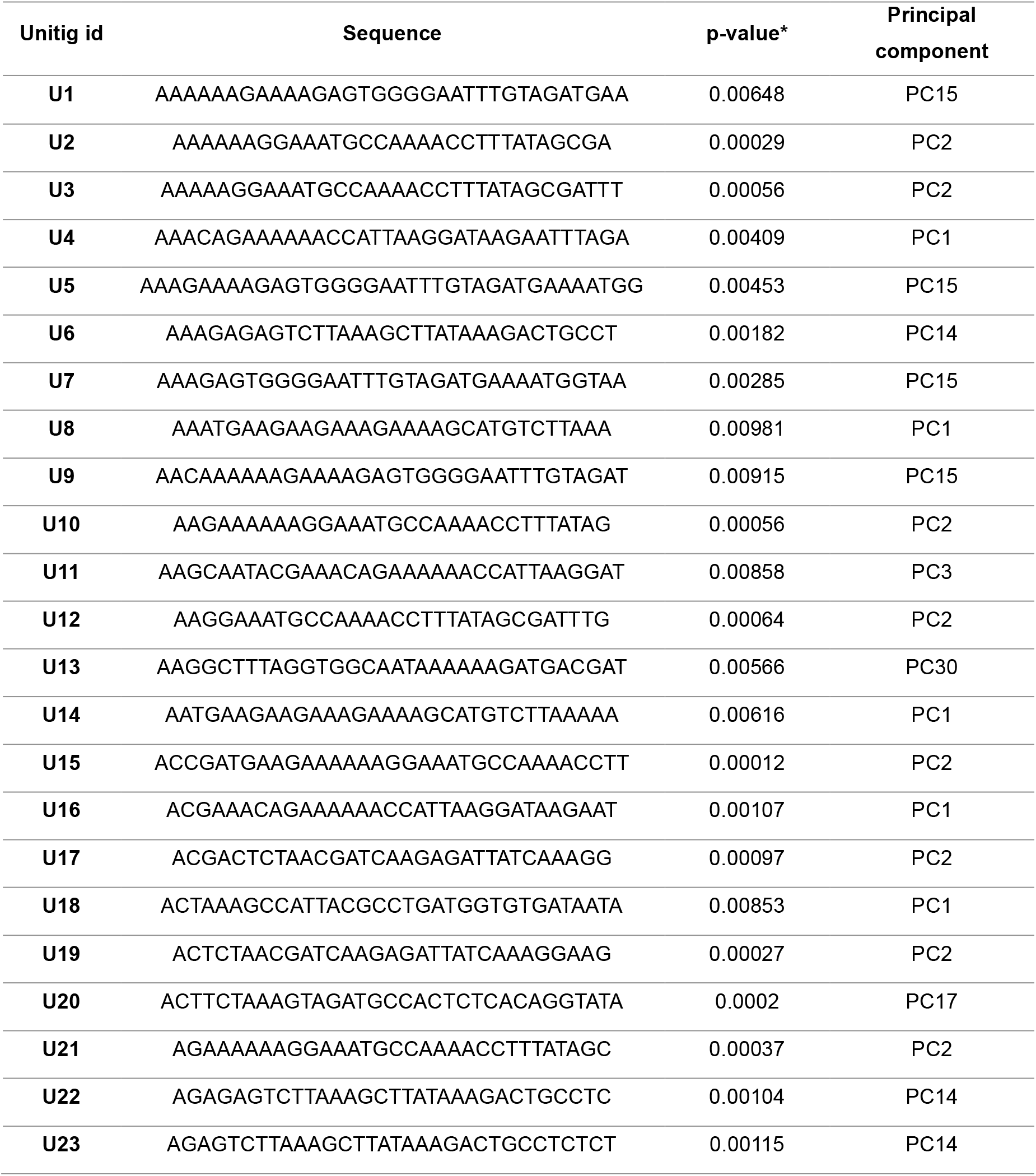
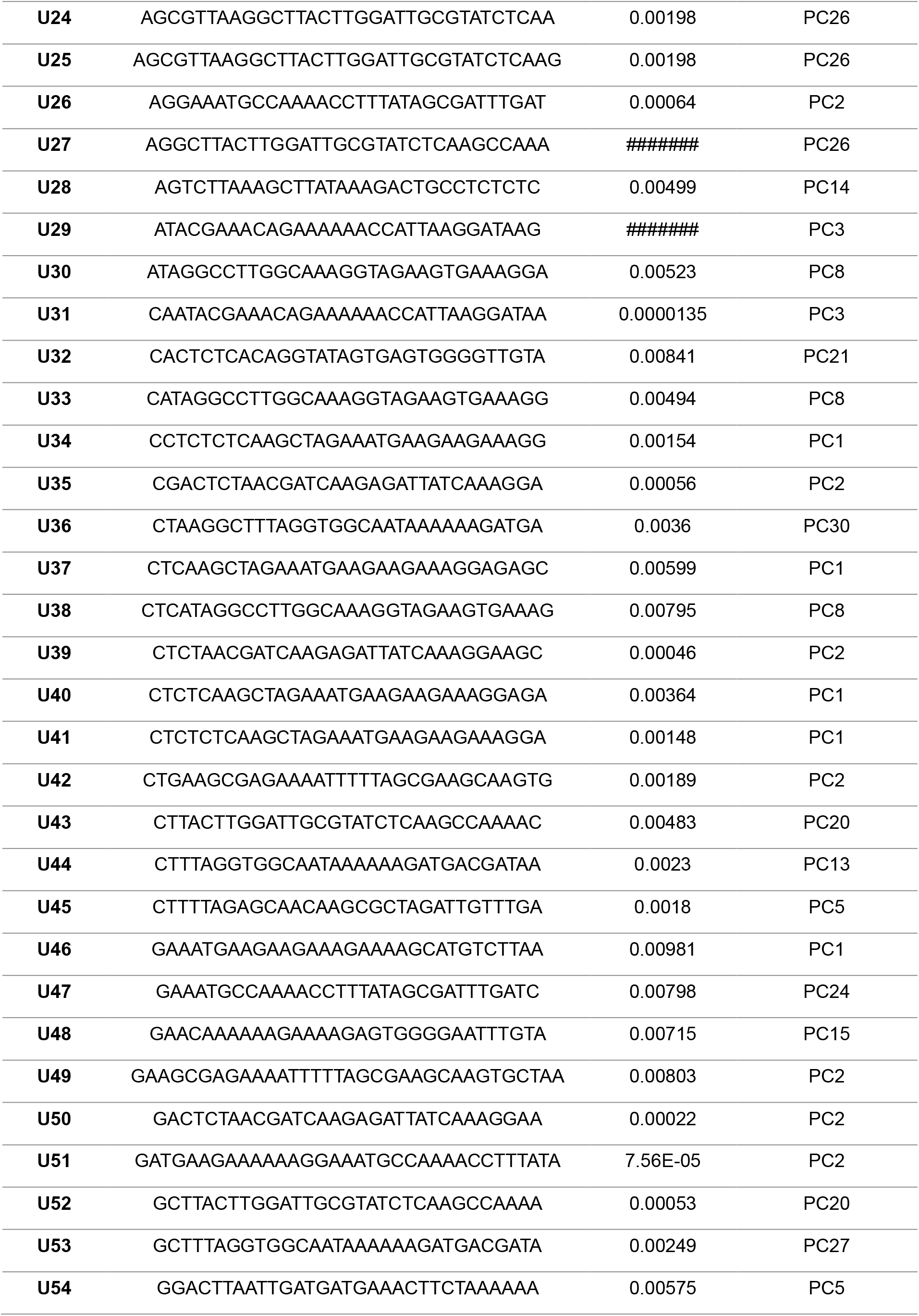
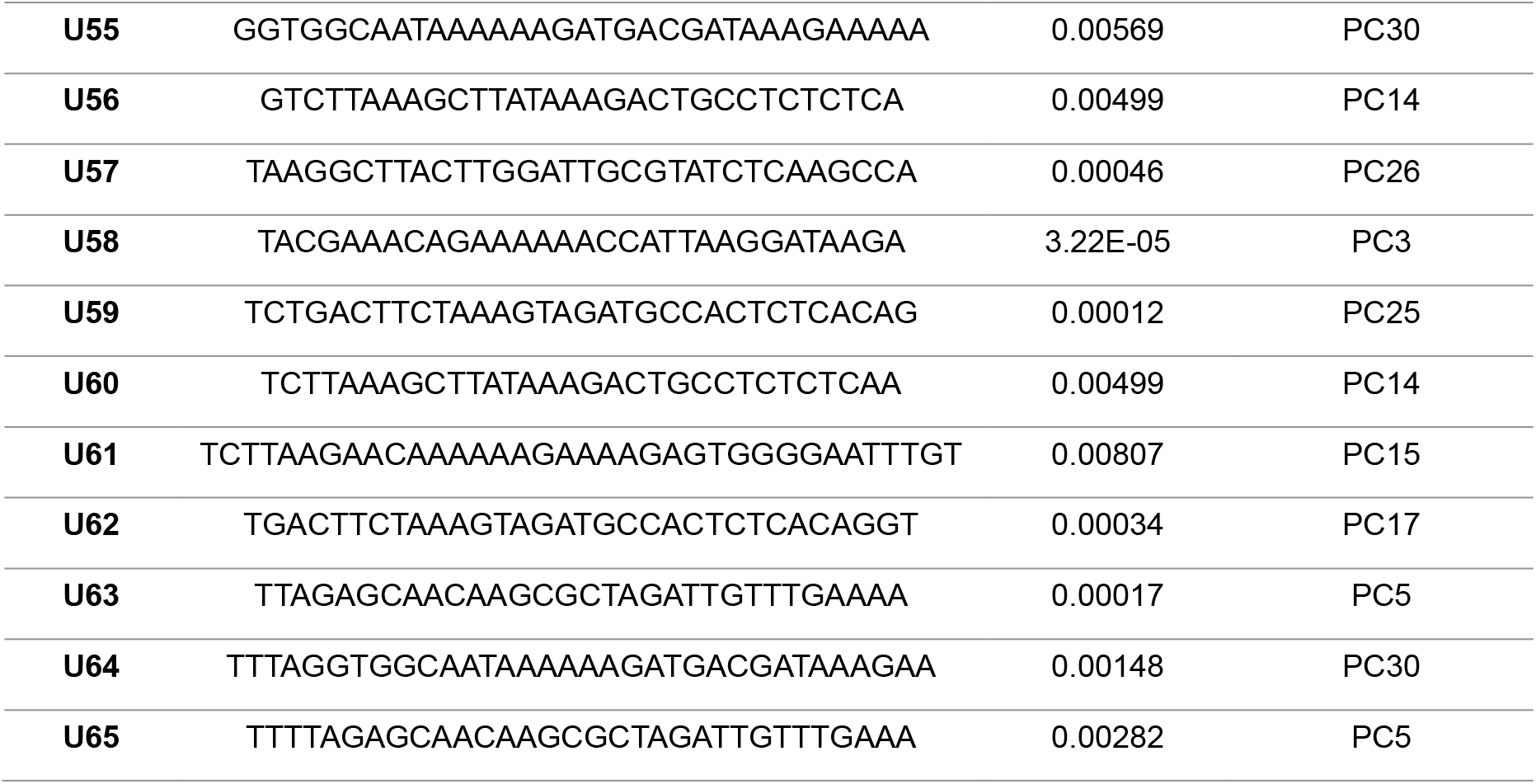
Unitigs significantly associated with GC (p-value) and its principal component.

**Supplementary Table 3.**
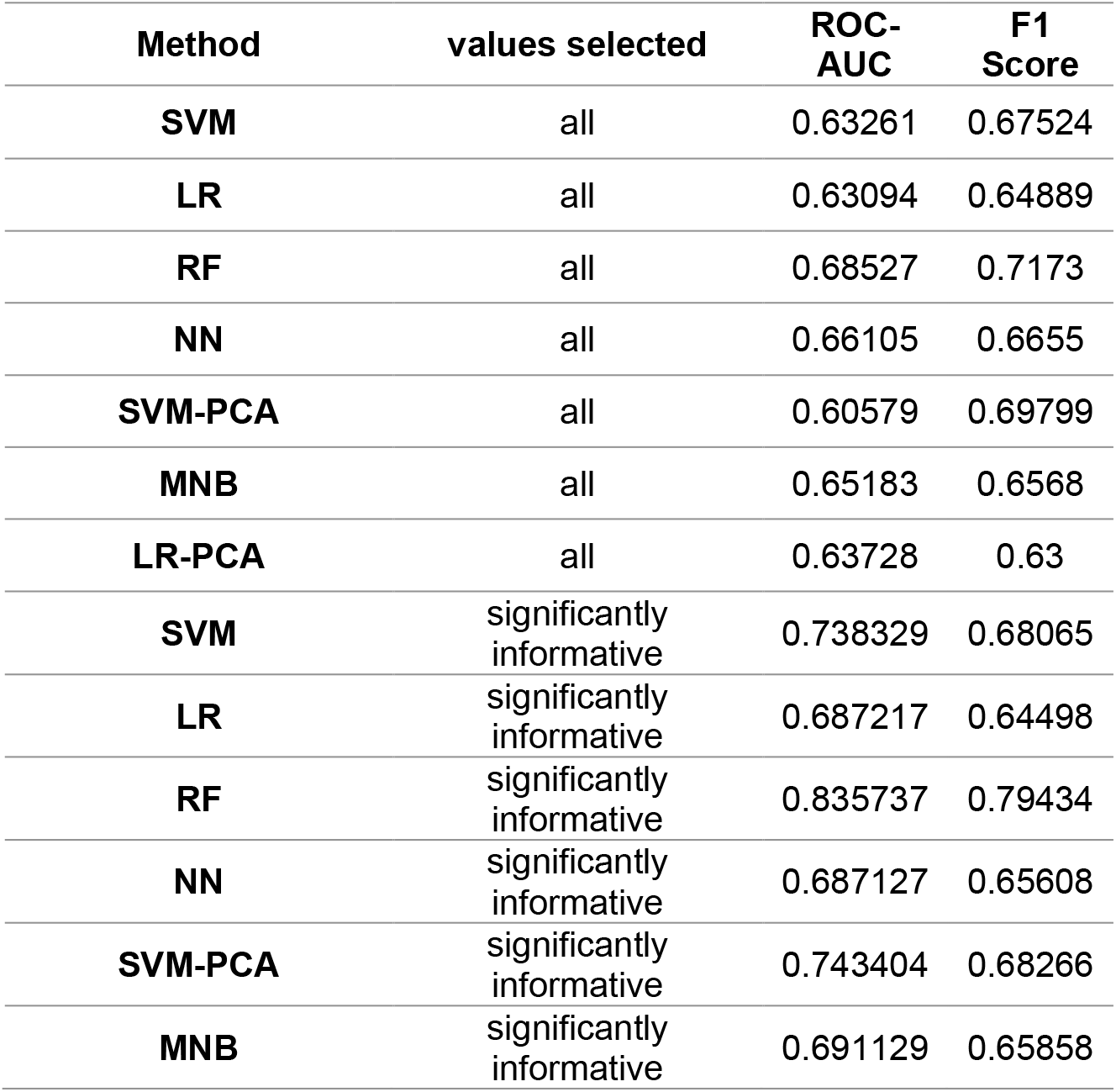
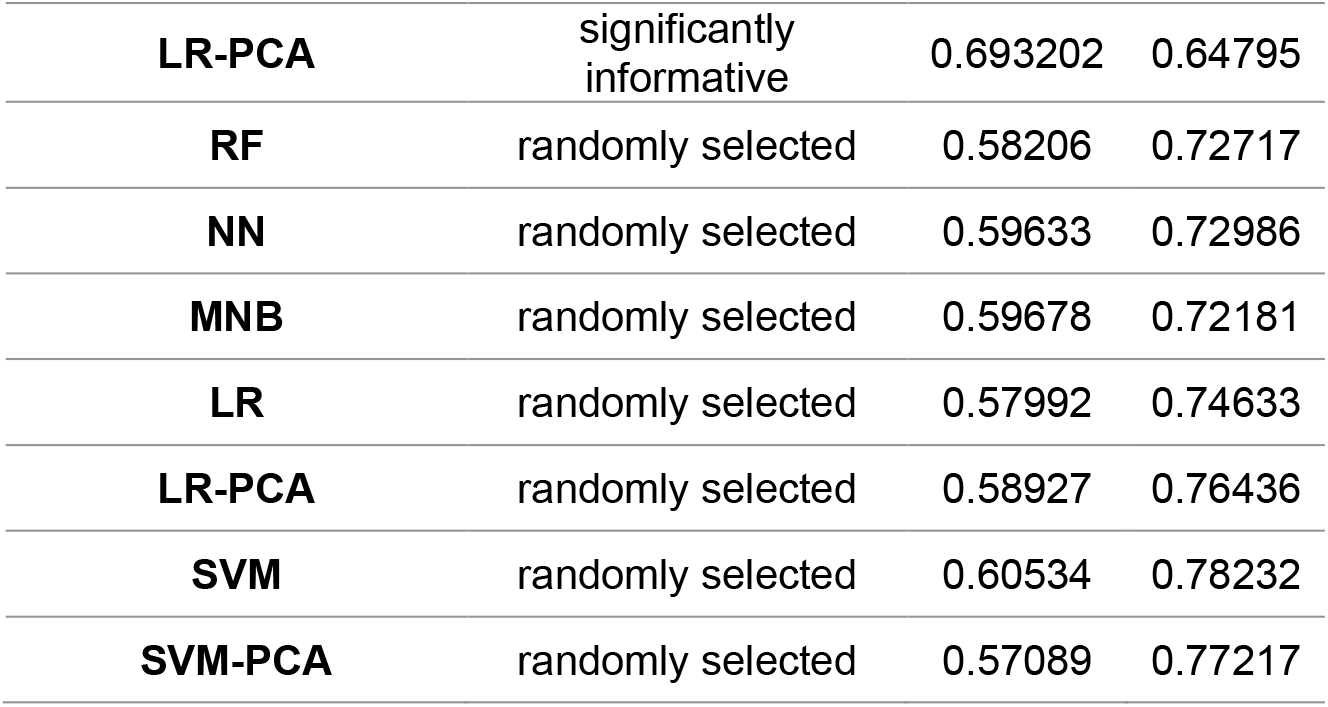
ROC-AUC and F1 scores of Machine Learning training for the six models tested, including all values, only the significantly informative and a set randomly choose

**Supplementary Table 4.**
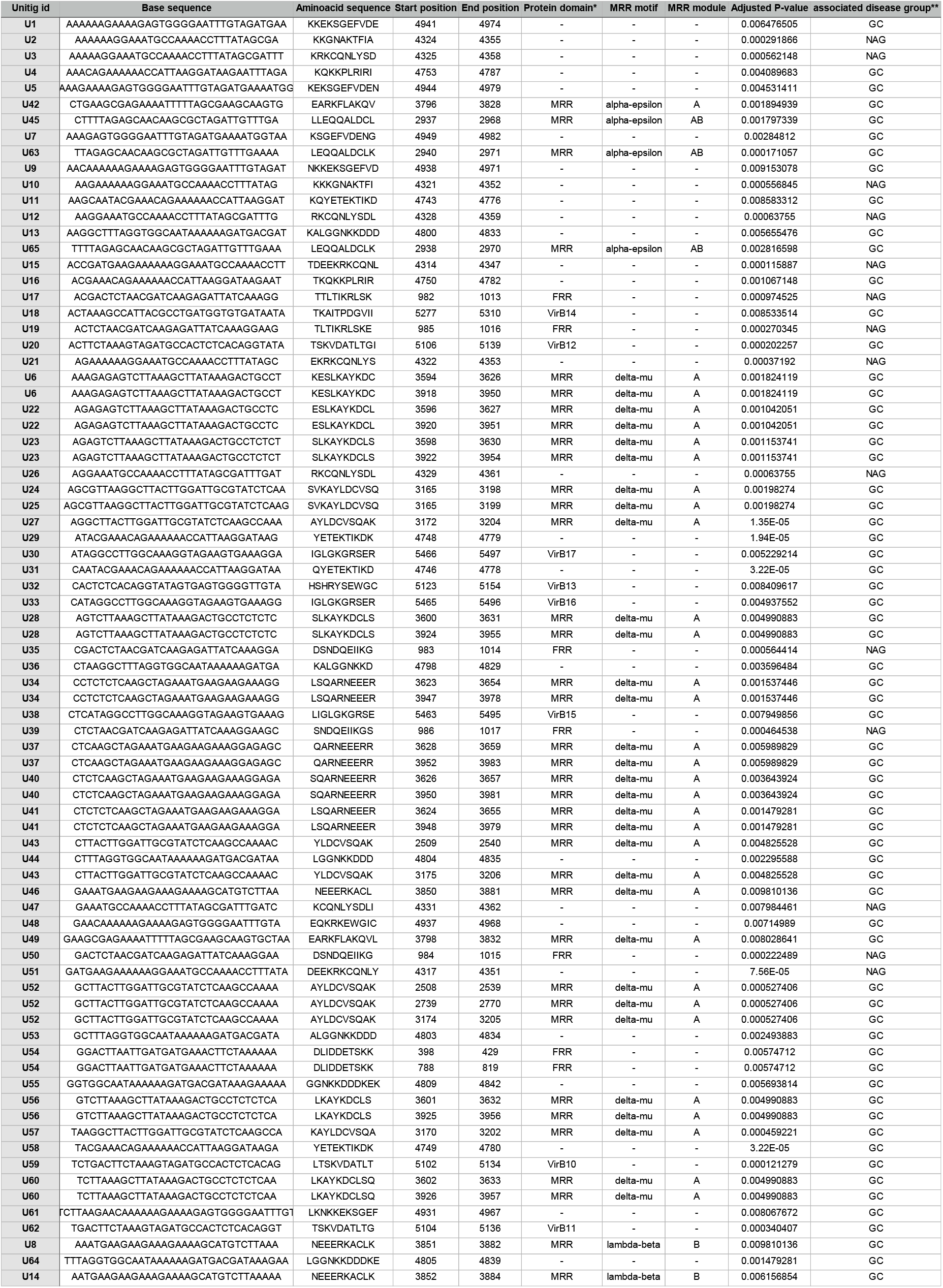
Position of the 89 significatively informative unitigs in the *cagY* gene of the reference strain HP-26695; p-value of the disease association

